# Zebrafish *Dscaml1* is Essential for Retinal Patterning and Function of Oculomotor Subcircuits

**DOI:** 10.1101/658161

**Authors:** Manxiu Ma, Alexandro D. Ramirez, Tong Wang, Rachel L. Roberts, Katherine E. Harmon, David Schoppik, Avirale Sharma, Christopher Kuang, Stephanie L. Goei, James A. Gagnon, Steve Zimmerman, Shengdar Q. Tsai, Deepak Reyon, J. Keith Joung, Emre R. F. Aksay, Alexander F. Schier, Y. Albert Pan

**Affiliations:** Department of Neuroscience and Regenerative Medicine, Medical College of Georgia, Augusta University, Augusta, GA 30912; Center for Neurobiology Research, Fralin Biomedical Research Institute at Virginia Tech Carilion, Virginia Tech, VA 24016; Institute for Computational Biomedicine and the Department of Physiology and Biophysics, Weill Cornell Medical College, New York, New York 10021; Graduate Program in Neuroscience, Augusta University; Depts. of Otolaryngology, Neuroscience & Physiology, and the Neuroscience Institute, New York University Langone School of Medicine, New York, NY 10016; Medical Scholars Program, Augusta University; Department of Ophthalmology, Medical College of Georgia, Augusta University, Augusta, GA 30912; Department of Molecular and Cellular Biology, Harvard Stem Cell Institute, Center for Brain Science, Harvard University, Cambridge, MA 02138; School of Biological Sciences, University of Utah, Salt Lake City, Utah 84412; Molecular Pathology Unit, Center for Computational and Integrative Biology, and Center for Cancer Research, Massachusetts General Hospital, Charlestown, MA 02129; Department of Pathology, Harvard Medical School, Boston, MA 02115; Department of Hematology, St. Jude Children’s Research Hospital, Memphis, TN, 38105; Editas Medicine, 11 Hurley Street, Cambridge, MA 02142; The Broad Institute of Massachusetts Institute of Technology and Harvard, Cambridge, MA 02142

**Author notes:** T.W., M.M., and A.R. contributed equally to this article. Correspondence should be addressed to Y. Albert Pan, Center for Neurobiology Research, Fralin Biomedical Research Institute at VTC, 2 Riverside Circle, Roanoke, VA 24016. **Conflict of Interest:** J.K.J. has financial interests in Beam Therapeutics, Editas Medicine, Pairwise Plants, Poseida Therapeutics, Transposagen Biopharmaceuticals, and Verve Therapeutics. J.K.J.’s interests were reviewed and are managed by Massachusetts General Hospital and Partners HealthCare in accordance with their conflict of interest policies. J.K.J. holds equity in Excelsior Genomics. J.K.J. is a member of the Board of Directors of the American Society of Gene and Cell Therapy. J.K.J. is a co-inventor on various patents and patent applications that describe gene editing and epigenetic editing technologies.

## Abstract

<u>D</u>own <u>S</u>yndrome <u>C</u>ell <u>A</u>dhesion <u>M</u>olecules (*dscam* and *dscaml1*) are essential regulators of neural circuit assembly, but their roles in vertebrate neural circuit function are still mostly unexplored. We investigated the role of *dscaml1* in the zebrafish oculomotor system, where behavior, circuit function, and neuronal activity can be precisely quantified. Loss of zebrafish *dscaml1* resulted in deficits in retinal patterning and light adaptation, consistent with its known roles in mammals. Oculomotor analyses showed that mutants have abnormal gaze stabilization, impaired fixation, disconjugation, and faster fatigue. Notably, the saccade and fatigue phenotypes in *dscaml1* mutants are reminiscent of human ocular motor apraxia, for which no animal model exists. Two-photon calcium imaging showed that loss of *dscaml1* leads to impairment in the saccadic premotor pathway but not the pretectum-vestibular premotor pathway, indicating a subcircuit requirement for *dscaml1*. Together, we show that *dscaml1* has both broad and specific roles in oculomotor circuit function, providing a new animal model to investigate the development of premotor pathways and their associated human ocular disorders.

## Introduction

The Down Syndrome Cell Adhesion Molecule (DSCAM) family proteins are neuronal cell recognition molecules that are essential for neural circuit assembly and neural patterning across different phyla. In humans, loss of DSCAM and DSCAM-like 1 (DSCAML1) are linked to autism spectrum disorder and cortical abnormalities^1,2^. At the cellular level, DSCAMs promote avoidance between neurites of the same cell or between similar cell types. DSCAMs are also involved in the regulation of cell death, synaptic adhesion, axon outgrowth, axonal refinement, and dendritic growth^3, 4, 5, 6, 7, 8^. However, it is still unclear how the sum of these cellular functions contribute to neural circuit activity, behavior, and human disorders.

Here, we investigate the role of DSCAML1 on neural circuit function in the context of oculomotor behavior, a setting with a rich history and easy to quantify behaviors. Eye movement is an integral part of visual perception and survival in vertebrates, enabling shifts in visual attention and preventing image blurring during object or head motion. We focus on saccade, fixation, and gaze stabilization movements, which are highly conserved among vertebrates. Saccades are fast ballistic eye movements that allow rapid gaze shifting and redirection of visual attention. After each saccade, gaze is maintained at a single location, termed fixation. Saccades also occur reflexively during gaze stabilization, *e.g*., the optokinetic reflex (OKR) and vestibular ocular reflex (VOR)^9^. OKR is triggered by broad-field visual motion, during which the eyes counteract motion by velocity-matched eye movements. The periods of velocity matching, called ‘slow phase’, are interrupted by ‘fast phase’ saccadic movements to the opposite direction, which reset eye position. Slow and fast phases are also triggered by the vestibular system in the VOR to counteract head motion.

In the oculomotor system, the neural circuitry is mapped in detail, and changes to eye movement kinematics can be attributed to specific subcircuit deficits. Sensory inputs and motor commands from higher brain centers converge onto the brainstem, where premotor neurons generate motor signals encoding eye position and velocity. The premotor pathways, in turn, activate the cranial motor neurons that innervate the extraocular muscles (Fig. 1A). Three subcircuits are the focus of this study. The saccadic premotor pathway, which goes through the excitatory burst neurons in the midbrain and hindbrain, initiates saccade and encodes the appropriate eye velocity. The vestibular premotor pathway, which receives input from the eye and ear, controls slower eye movement and velocity matching during gaze stabilization. Lastly, the neural integrator premotor pathway integrates information from the saccade and vestibular pathways to maintain stable eye position over time. Deficits in these subcircuits are associated with human diseases such as ocular motor apraxia (saccade deficit), optokinetic response abnormalities (vestibular deficit), and gaze-evoked nystagmus (neural integrator deficit)^9^. Eye movement deficits are common in the general population and can lead to severely impaired visual function and difficulty in daily tasks such as reading or crossing the street. Saccade deficits are also highly prevalent in neuropsychiatric and neurodegenerative patient populations, serving as a diagnostic tool and a window into the underlying pathophysiology^10, 11^. However, our understanding of the genetic contributors for oculomotor circuit development and neural mechanisms for eye movement disorders are still limited.

**Figure 1.**
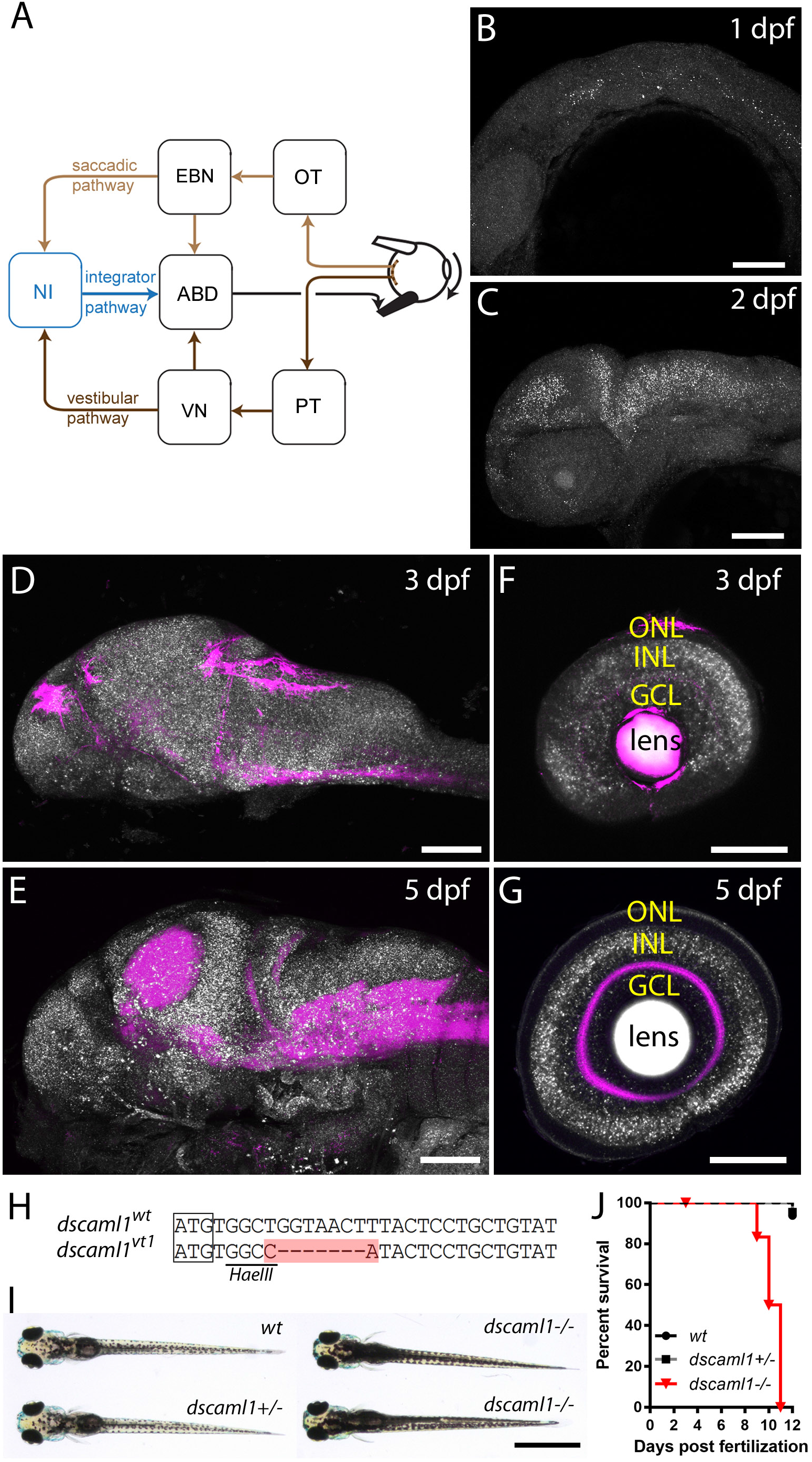
The Oculomotor circuit, and the expression and gene targeting of *dscaml1*. **A**, Diagram of the oculomotor circuit for horizontal eye movement. Visual motion activates directional selective retinal ganglion cells, which innervate the optic tectum (OT) and pretectum (PT). PT provides input to the vestibular nucleus (VN). OT (along with other areas), activates the excitatory burst neurons (EBN). EBN and VN provide premotor input to the abducens nucleus (ABD), which innervates the extraocular muscles. EBN and VN also innervate the velocity-position neural integrator (NI), which provides an eye position signal to the ABD. The saccadic, vestibular, and integrator pathways are labeled light brown, dark brown, and blue, respectively. **B-G**, *dscaml1 mRNA* (white) labeled by fluorescent *in situ* hybridization, with neuropil counterstained with an antibody against synaptotagmin 2 (Znp-1, magenta). Developmental staging as indicated. **H**, Alignment of wild-type and TALEN targeted *dscaml1* genomic sequence. The start codon is boxed, and the region containing insertions and deletions is highlighted in red. **I**, Pigmentation pattern of different genotypes. *dscaml1* mutant animals show darker overall pigmentation. **J**, Survival curve, sorted by genotype. GCL: ganglion cell layer; INL: inner nuclear layer; ONL: outer nuclear layer. Scale bar is 100μm in panel A-F, 1mm in H.

We take advantage of the zebrafish oculomotor system as a framework to probe the role of *dscaml1*, the zebrafish homolog of *DSCAML1*, in neural circuit development, neuronal activity, and behavior^12^. Zebrafish oculomotor behavior is robust at the larval stage; the small size and optically translucency of zebrafish larvae make it possible to use two-photon calcium imaging to record neuronal activity *en masse* while the animal is behaving^13^. Additionally, the ontogeny and anatomy of the fish oculomotor system is well characterized and conserved with that of mammals^14,15^. We found that genetic deletion of zebrafish *dscaml1* affected retinal patterning, light adaptation, and oculomotor behaviors. Our results suggest that for horizontal eye movements, the saccade and integrator subcircuits are affected in *dscaml1* mutants, while the vestibular subcircuit is mostly spared. *dscaml1* also affects the robustness of OKR against prolonged visual stimulation. The mutant phenotype mirrors several characteristic features of congenital ocular motor apraxia (COMA) (OMIM 257550) and provides insight into the potential neural mechanisms of COMA and other human oculomotor disorders^16,17^.

## Results

To investigate the role of *dscaml1* in visual circuit activity and function, we first examined its expression in zebrafish. Then we assessed how the loss of *dscaml1* affected visual pathway development and locomotor behavior. Finally, we analyzed oculomotor behavior and neural activity to deduce the underlying neural circuit changes in *dscaml1* mutants.

### Expression and gene targeting of zebrafish *dscaml1*

We examined the expression of *dscaml1* with whole-mount fluorescent *in situ* hybridization, from the embryonic stage to five days post-fertilization (dpf), during which visually guided eye movements begin to mature^18,19^. *dscaml1* expression was enriched in the nervous system and present in most brain regions (Fig. 1B-E). In the retina, *dscaml1* was expressed in the inner nuclear layer (INL, containing the amacrine, bipolar, and horizontal cells) and the ganglion cell layer (GCL, containing retinal ganglion cells and amacrine cells) at 3 dpf (Fig. 1F-G). At 5 dpf, *dscaml1* expression was seen predominantly in the INL and sparsely in the GCL. These early and late expression patterns match the expression of mouse *Dscaml1* in the retina at P6 and P12, respectively^12,20^.

To test the function of *dscaml1*, we generated a mutant allele of *dscaml1* by <u>TAL</u>-like effector nucleases (TALENs) mediated gene targeting^21^. The mutant allele, *dscaml1^vt1^*, contains a seven base pair deletion (9 base pair deletion plus 2 base pair insertion) after the start codon (Fig. 1H). Incorporation of the TALEN-mediated deletion in the *dscaml1* mRNA was confirmed by RT-PCR and sequencing. The deletion results in frame shift and a truncated open reading frame, which lacks the signal peptide and all functional domains. These results suggest *dscaml1^vt1^* is likely a null allele.

Homozygous mutant animals showed behavioral and morphological changes associated with the visual system. At 5 dpf, homozygous mutant animals (*dscaml1−/−*) appeared to develop normally, with normal head size, normal eye size, and inflated swim bladders. Compared with heterozygous and wild-type siblings, however, mutant animals had darker pigmentation when placed on a light background (Fig. 1I). This deficit in background adaptation, a camouflage response that requires light detection, suggests that *dscaml1−/−* animals may be visually impaired^22^. At later stages, lethality was observed in mutant animals at 8 dpf, and none survived past 11 dpf (Fig. 1J). Heterozygous animals *(dscaml1+/−)* are viable and fertile as adults. Despite the late larval stage lethality, *dscaml1* mutant brain showed no gross abnormalities. Major longitudinal and commissural axon tracts, motor axon terminals, and peripheral ganglia are formed normally in mutants (Supplementary Fig. S1). The only observed morphometric difference was in the optic tectum neuropil region, a major retinorecipient area important for sensory processing. The optic tectum neuropil was 23% larger in *dscaml1* mutants (44,121±1,017 μm^2^) versus wild type (35,816±656.5 μm^2^) or heterozygotes (36,715±549 μm^2^) (p<0.0001, one-way ANOVA) (Supplementary Fig. S1D).

### *dscaml1* is required for planar and laminar patterning in the retina

Genetic studies in mice and chick demonstrated that Dscaml1 is required for planar patterning of retinal amacrine cells and laminar specific neurite termination in the inner plexiform layer (IPL)^6, 20, 23^. We found that *dscaml1* has conserved functions in retinal patterning in zebrafish (Fig. 2). With H&E staining, we did not see visible perturbation of retina structure, with the exception that the IPL is thicker in the heterozygotes and mutants, compared to wild types (Fig 2A-B). Increased IPL thickness was also seen in *Dscam* and *Dscaml1* mutant mice, and may be caused by decreased developmental cell death^20,23^. To test *dscaml1’s* role in planar patterning, we labeled the serotonin (5-HT) expressing amacrine cells. 5-HT cells represent a single amacrine cell type (S1) that is sparsely distributed in the retina, with very few contacts between cell bodies (Fig. 2C)^24^. In *dscaml1* mutants, however, frequent clustering of 5-HT amacrine cells was observed. We calculated the probability of one cell being immediately adjacent to another, termed aggregation index (density recovery profile analysis was not possible due to the sparsity of this cell type in the larvae). Compared to wild-type animals, the aggregation index in mutants was significantly increased (Fig. 2D). The increased aggregation seen in the mutants was not due to an increase in the number of 5-HT amacrine cells, which was not significantly different between wild-type, heterozygote, and mutant animals (Fig. 2E).

**Figure 2.**
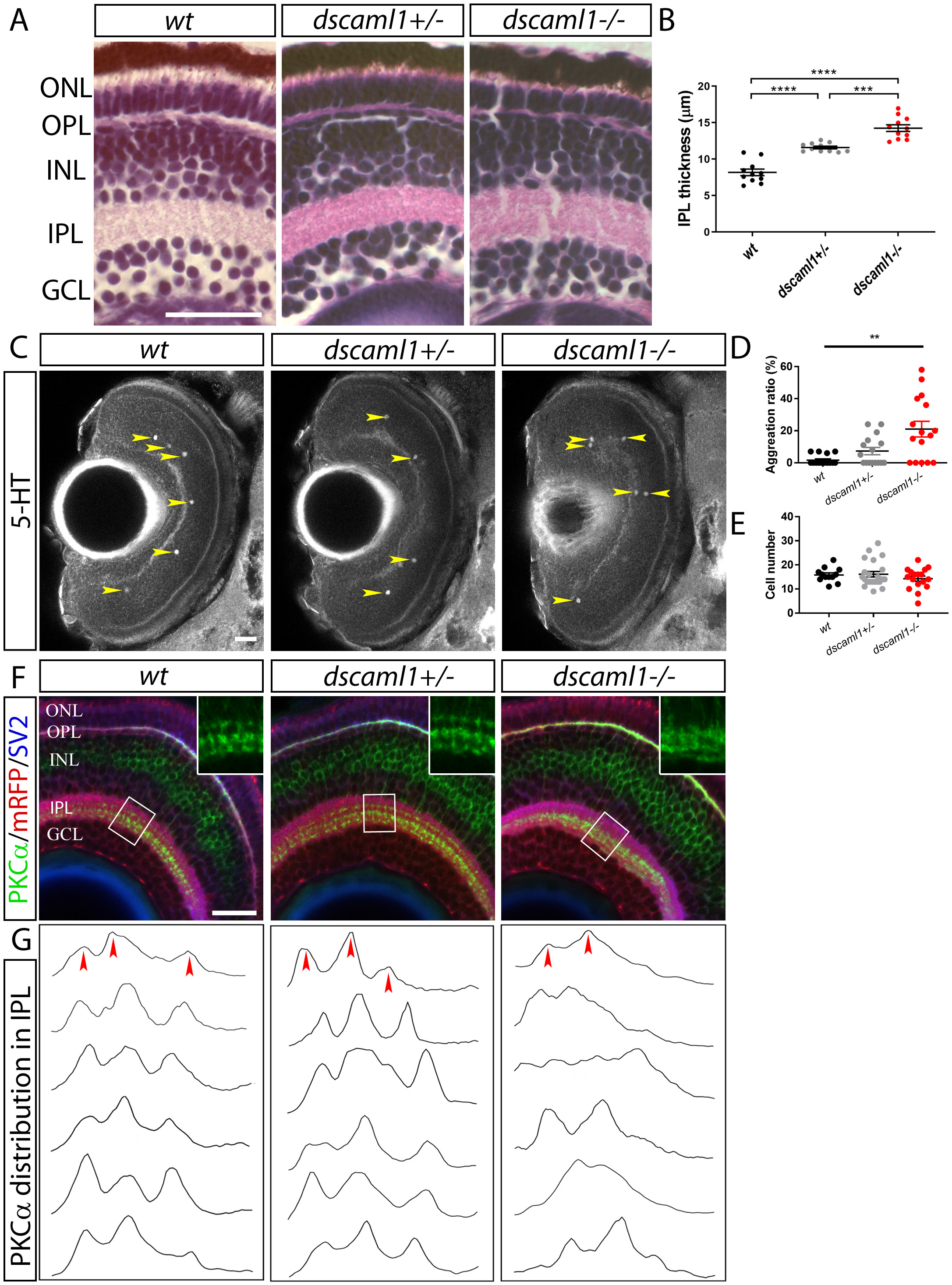
*dscaml1* is required for planar and laminar patterning of the retina. **A**, H&E staining of 5 dpf retina. **B**, Quantification of IPL thickness, performed in confocal imaged 5 dpf larvae immunostained with Znp-1 antibody. Mutants have significantly thicker IPL compared to wild-type animals. Heterozygotes have an intermediate phenotype. **C**, Serotonergic amacrine cells, stained with an antibody against 5-hydroxytryptamine (5-HT). Cell bodies are indicated by yellow arrowheads. **D**, Percentage of cells that are immediately adjacent to another cell (within 10μm, approximately 3 cell diameter) is significantly higher in *dscaml1−/−* animals (**p<0.01, one-way ANOVA). **E**, The number of 5-HT positive amacrine cells was not significantly different. **F**, Immunostaining of ON-bipolar cell (PKCα, green), retinal ganglion cell *[Tg(atoh7:GAP-RFP)*, red], and synapses (SV2, blue). Boxed areas are enlarged two-fold and shown in the inset. Both mutants and morphants show loss of discrete stratification and mature terminal boutons in the ON (lower) sublamina of the IPL. **G**, PKCα immunolabeling intensity (vertical axis) was plotted across the depth of the IPL (horizontal axis), normalized for maximal intensity and IPL thickness. Three prominent peaks can be discerned in wild-type and heterozygote animals (red arrowheads). The number and location of peaks are more variable in mutants and morphants. GCL: ganglion cell layer; INL: inner nuclear layer; IPL: inner plexiform layer; ONL: outer nuclear layer; OPL: outer plexiform layer. Scale bars are 25μm. One-way ANOVA was used for statistical comparisons in D and E (****p<0.0001, ***p<0.001, **p<0.01).

We also found that loss of *dscaml1* impacted laminar specific neurite termination in the IPL, specifically of the ON-bipolar cell axon terminals (visualized with anti-PKCα)^25^. ON-bipolar cells transduce electrical activity in response to light increments (e.g., lights turning on) and have dendrites that extend toward the OPL (where they synapse with photoreceptor cells) and axons that form three discrete layers in the inner half of the IPL (the ON sublamina) (Fig. 2F)^26^. This distribution was quantified by measuring the fluorescent intensity of PKCα immunolabeling across the thickness of the IPL, as defined by SV2 immunolabeling^27^. In all wild-type and heterozygote animals, three distinct peaks at stereotypical positions can be discerned. In contrast, PKCα distribution in the mutants was more diffuse and did not form three discrete sublaminae (Fig. 2G). Cone photoreceptors and Müller glia appear morphologically normal in the *dscaml1* mutant retina (Fig. 3A-F).

**Figure 3.**
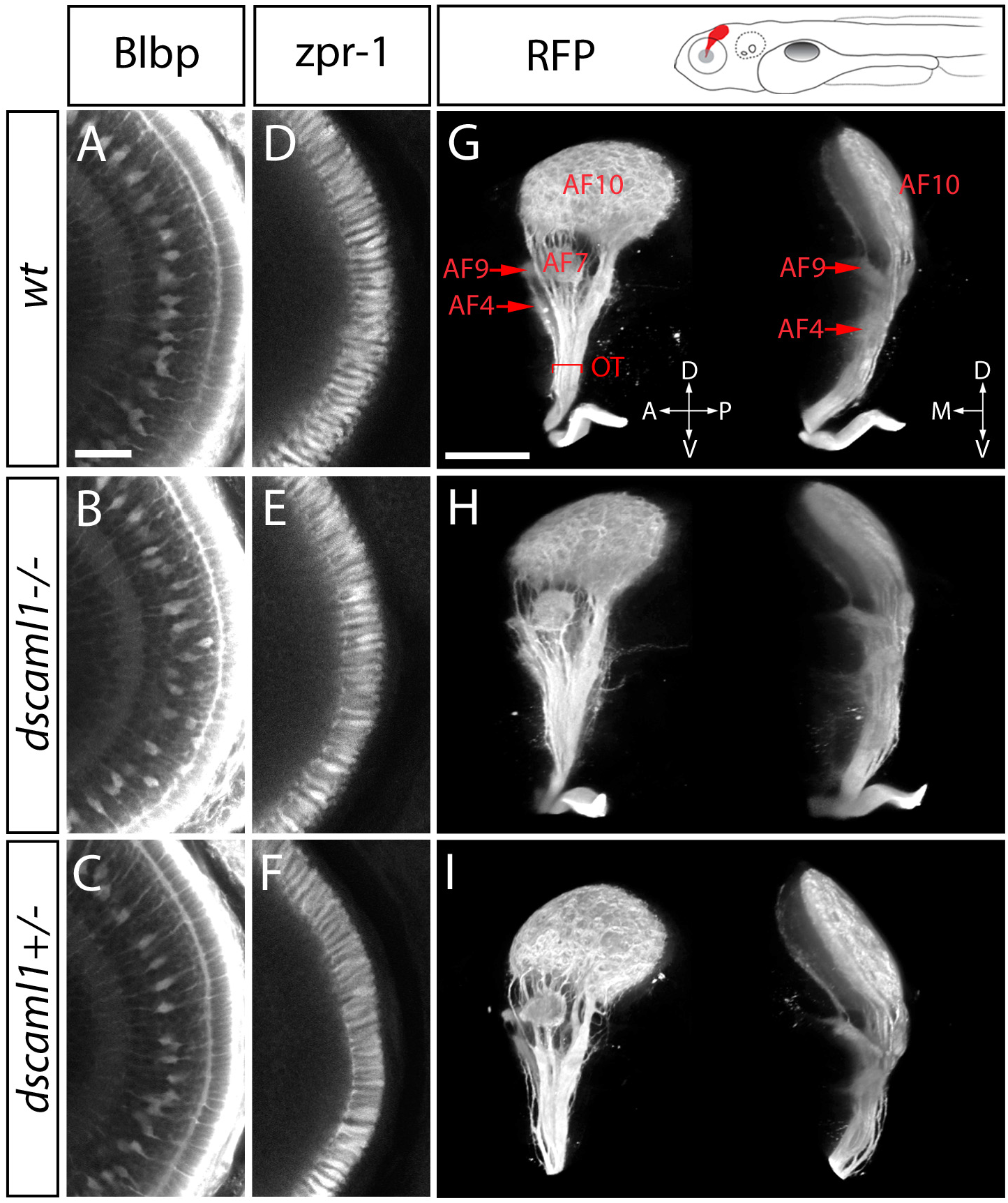
Normal development of Müller glia, outer retina, and optic tract in the *dscaml1* mutant. **A-F**, No abnormalities were seen in cone photoreceptor cells (stained with zpr-1 antibody) and Müller glia (anti-Blbp). **G-I**, Retinal afferent projection in *dscaml1* mutants. The panel above G shows an illustration of retinal afferents (red) in the *Tg(atoh7:GAP-RFP)* transgenic. Images show the 3D reconstruction of retinal afferents viewed from the side (left images) and front (right images). Retinal arborization fields (AFs) in the thalamus (AF4), pretectum (AF7, 9), and optic tectum (AF10) are identified as described previously^29^. No differences were observed among different genotypes. OT: optic tract. Panels A-F are shown at the same scale, scale bar in A is 25 μm. Panels G-I are shown at the same scale, scale bar in G is 100 μm.

Loss of *dscaml1* did not appreciably affect the afferent projections of the retina. We utilized the *Tg(atoh7:GAP-RFP)* transgenic line, which expresses membrane-tagged red fluorescent protein in retinal ganglion cells^28^. In *dscaml1* mutants, the optic nerve crossed the chiasm normally, and the axon terminals (arborization fields) of the optic tract were mostly indistinguishable from wild-type and heterozygous animals (Fig. 3G-I)^29^. This suggests that the expression of *dscaml1* in the GCL is not required for outgrowth of retinal ganglion cell axons. It remains possible, however, that the projections of a subset of retinal ganglion cells may be affected by the loss of *dscaml1*. Overall, the expression and mutant phenotypes of zebrafish *dscaml1* demonstrate that it is functionally conserved with other vertebrates and required for both planar and laminar patterning in the retina.

### Abnormal locomotor behavior in *dscaml1* mutants

Given the retinal patterning deficits and background adaptation abnormality seen in the mutants, we asked whether light-induced locomotor activity is also affected. We monitored locomotor activity of individual 5 dpf larvae over 24 hours, under normal circadian cycle (14 h day/10 h night) (Fig. 4A)^30,31^. Consistent with previous studies, wild-type fish were more active during the day, less active at night, and responded robustly during day/night and night/day transitions^30^. This diurnal rhythm was preserved in *dscaml1* mutants. Loss of *dscaml1* did not affect the duration of active periods, during either day or night (Fig. 4B, C). The amount of movement, however, was reduced in mutants during the day as well as the response to lights turning on, compared to wild-type and heterozygous animals (Fig. 4D-F). Interestingly, the response in mutants to lights off was as robust as for wild-type and heterozygous siblings (Fig. 4G). These results show that *dscaml1* mutants can detect light and move in response to a light offset, but are deficient in responding to light onset. These behavioral deficits are consistent with a partial deficit in the retinal ON pathway, potentially contributed by imprecise sublamina-specific ON-bipolar cell axon targeting.

**Figure 4.**
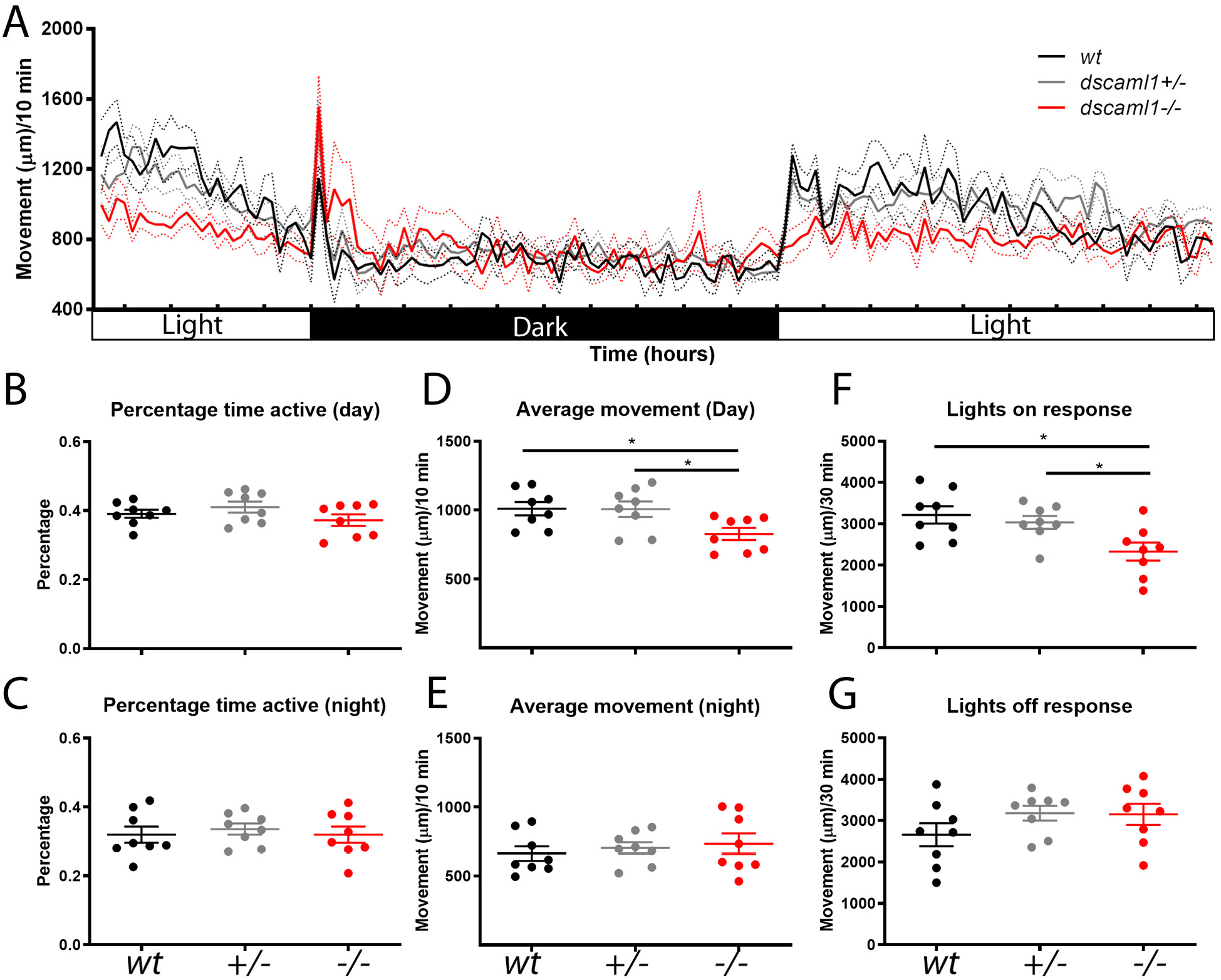
Locomotor activity in response to light. **A**, Locomotor activity over 24 hours. Solid lines are mean movement (n=8 for each group), and dotted lines indicate the range of standard error. Lighting conditions are indicated on the X-axis, with each tick marking one hour. **B-C**, Percentage time active during the day (B) and night (C). **D-E**, Average movement during the day (D) and night (E). **F-G**, Total amount of movement 30 minutes after lights switch on (F) and off (G). *p<0.05 for pairwise comparisons using one-way ANOVA.

### Abnormal OKR and eye lock up in *dscaml1* mutants

To further test the role of *dscaml1* in neural circuit function and sensorimotor integration, we examined the mutants’ performance in OKR. OKR consists of visual motion (i.e., optic flow)-triggered tracking eye movements (slow phases) and intervening resetting saccades (fast phases) (Fig. 5A). The quality of OKR is measured by slow-phase gain, which is the ratio of eye velocity to the velocity of the visual motion. In zebrafish, OKR develops early and is robust by 3-4 dpf^18^. We tested 5-6 dpf larvae inside a circular arena where black and white moving gratings were projected onto the arena, and eye position was video-recorded simultaneously (Fig. 5A, Movie 1). Grating directions alternated between clockwise and counter-clockwise at 3 or 40 seconds (Fig. 5B, E). Under short time scale (3 s), fast phase has little effect on OKR performance, and gain is directly related to the processing of optic flow via the vestibular premotor pathway (Fig. 1A)^32,33^. Under long time scale (40 s), saccades are necessary to reset eye position periodically, and the oculomotor system needs to be robust against eye position drifts and fatigue in optic flow response^19,34,35,36^.

**Figure 5.**
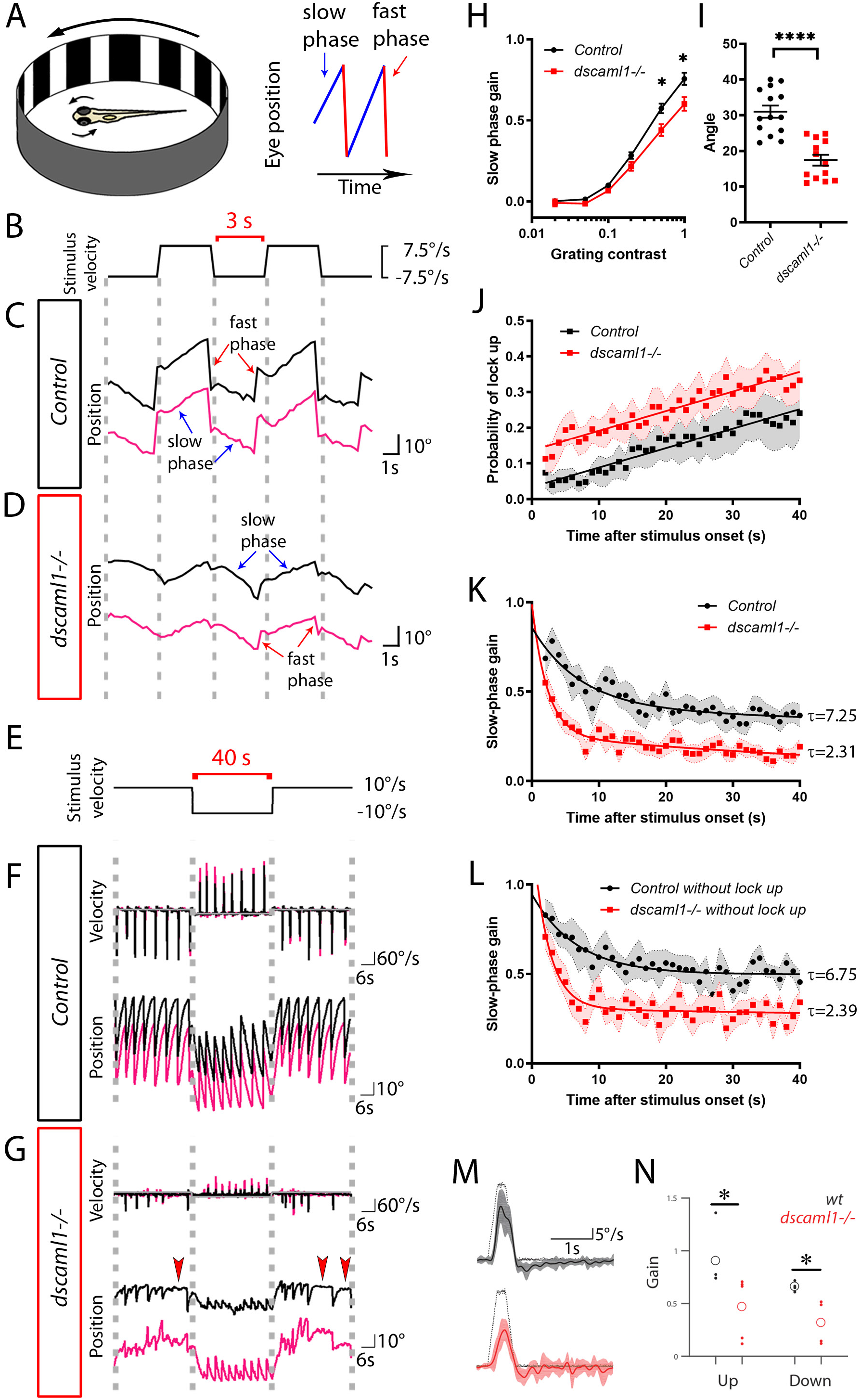
OKR and VOR performance in *dscaml1* mutants. **A**, larvae were immobilized in the center of a circular arena, where black and white vertical bars are projected. Diagram of eye position is shown on the right, with fast phases (red segments) and slow phases (blue segments). **B-E**, short time-scale OKR. Eye position traces are shown for control (C) and *dscaml1−/−* (D) animals, at the same time scale as the square wave stimulus (B). Right and left eye traces are in black and fuchsia, respectively. E-G, long time-scale OKR. Eye velocity and position traces are shown for control (C) and *dscaml1−/−* (D), at the same time scale as the stimulus (E). In mutants, the eyes intermittently became locked up (red arrowheads). **H**, slow-phase velocity across a range of contrast levels. (*p<0.05, two-way ANOVA with Bonferroni correction). **I**, saccade amplitude is lower in mutants (****p<0.0001, two-sample t-test). **J-K**, the temporal dependency of lock-up probability (J), slow-phase gain (K), and slow-phase gain excluding lock-up periods (L). Positive gains in H, J-K are defined as eye movement in the same direction as the stimulus. Regression lines, means, and standard error (shaded areas) are shown. Linear regression model was used in J and control group in K. Two phase exponential decay regression model with plateau constrained to zero was used for *dscaml1−/−* group in K and both groups in L. **M**, eye velocity in response to a 35°/s tilt stimulus (black dotted line) in wild-type (top panel, solid black line) and *dscaml1* mutant (bottom panel, solid red line). Shaded areas indicate interquartile range. Torsional velocity was slower in mutants, compared to wild type. **N**, velocity gain of upward and downward tilts, both of which are lower in mutants. Mean values are marked by open circles. *p<0.05, Mann-Whitney *U* test.

We found that visuomotor processing for optic flow is intact in *dscaml1* mutants. Under short time scale, both control (wild type and heterozygote) and mutant animals exhibited qualitatively normal optokinetic responses (Fig. 5B-D). Slow-phase gain increased linearly with the logarithm of stimulus contrast in both groups, and there was no difference in gain at lower contrast levels (contrast<0.2) (Fig. 5H). Mutants do have lower slow-phase gain than the controls at higher contrast levels (contrast=0.5 or 1), though the deficit is mild. These findings suggest that *dscaml1* is likely not required for the assembly or function of the vestibular premotor pathway, but affects OKR performance at conditions that elicit higher eye velocity^32^.

Under long time scale, *dscaml1* mutants showed deficits in saccadic eye movements and exhibited more pronounced behavioral fatigue. In control animals, fast phase (resetting saccade) and slow phase alternated regularly, and saccades had consistent frequency and amplitude (Fig. 5F). In mutants, resetting saccades were irregular and reduced in amplitude (Fig. 5G, I). Interestingly, mutant eye movements would frequently pause, similar to the “lock up” phenotype seen in human with saccade deficits^16^. Lock-up periods tended to occur towards the end of the stimulus period, and the probability of lock up increased linearly over time, both in controls and mutants (Fig. 5J). The probability of lock up (time bins where velocity <1°/s) was significantly higher in mutants, compared to controls (comparison of linear regression intercepts, p<0.0001)^35,36^. Lock-up events could occur throughout the eye position range in the mutants, suggesting that lock up is not due to a failure to initiate faste phase at more eccentric positions (Supplementary Figure S2). Concurrently, relative to controls, mutants had more pronounced decay of slow-phase gain over time, even when lock-up periods were excluded (Fig. 5K, L)^19,37^. In both control and mutant data sets, the decay of slow-phase gain was well fit with a double exponential. Loss of *dscaml1* resulted in a drop in the contribution by the longer (later) component, and faster decay in the shorter (earlier) component (Table I., decay constant for short component shown in Fig. 5K, L). These results suggest that *dscaml1* has important roles in saccade generation and velocity maintenance.

**Table 1.** Double exponential decay fit for slow phase gain.

| With lock up | Control | *dscaml1*<sup>-/-</sup> |
| --- | --- | --- |
| Y<sub>0</sub> | 0.86 | 0.99 |
| Contribution of long component | 47.31 | 25.87 |
| Tau (short) | 7.25 | 2.31 |
| Tau (long) | 305.00 | 72.01 |

Table I. Double exponential decay fit for slow phase gain.
| Without lock up | Control | *dscaml1*<sup>-/-</sup> |
| --- | --- | --- |
| Y<sub>0</sub> | 0.9461 | 1.297 |
| Contribution of long component | 52.66 | 23.65 |
| Tau (short) | 6.749 | 2.385 |
| Tau (long) | 2.89E+08 | 472.4 |

### *dscaml1* is required for torsional eye movements during VOR

Our OKR assays tested the capacity for horizontal gaze stabilization, which requires the lateral and medial recti. To test the functionality of the other four extraocular motoneuron populations, we measured the gravito-inertial VOR in the torsional plane (Fig. 5M, N). In response to nose-up pitch tilts, larvae will use their superior oblique and inferior rectus muscles to counter-rotate the eyes. Similarly, following nose-down pitch tilts, larvae stabilize their gaze using the superior rectus and inferior oblique muscles^38^. As expected, wild-type larvae at 5dpf showed strong VOR in response to 15° steps up or down away from horizontal, with a slightly stronger response to nose-up tilts. *dscaml1* mutants were able to initiate VOR in a directionally appropriate manner but had lower gain (max eye velocity/max table velocity, 35°/s) to both stimuli. As fish develop VOR in the absence of light, this deficit is likely not due to any visual impairment^18^. We conclude that *dscaml1* is involved in the performance of both the OKR and VOR, and in both the horizontal and torsional planes.

### *dscaml1* is required for saccade and fixation

Given the pronounced reduction in fast-phase response and faster decay of slow-phase response during OKR, we next examined spontaneous eye movements in the absence of structured visual or vestibular stimulus to assess specific deficits in saccade generation and neural integrator function. Spontaneous eye movements in zebrafish consist of intermittent saccades, followed by periods of fixation, similar to mammalian scanning saccades^19, 34^. This behavior is used to direct visual attention to the temporal retina, where photoreceptor density is highest^39,40^. Spontaneous saccades are conjugated and usually alternate in direction, with typical angular velocity (sampled at 5 Hz) greater than 100°/s (Fig. 6A). In control animals, spontaneous saccades with velocity greater than 100°/s were the predominant type of eye movement (59%) (Fig. 6C, D). In mutant animals, high-velocity saccades were nearly absent (Fig. 6B-D). Instead of alternating in directions, sequential saccades often moved in the same direction, suggesting that these eye movements may be short of their intended target (hypometric). Saccades in mutants were also significantly more disconjugated, indicating that saccade initiation was bilaterally desynchronized (Fig. 6B, E).

**Figure 6.**
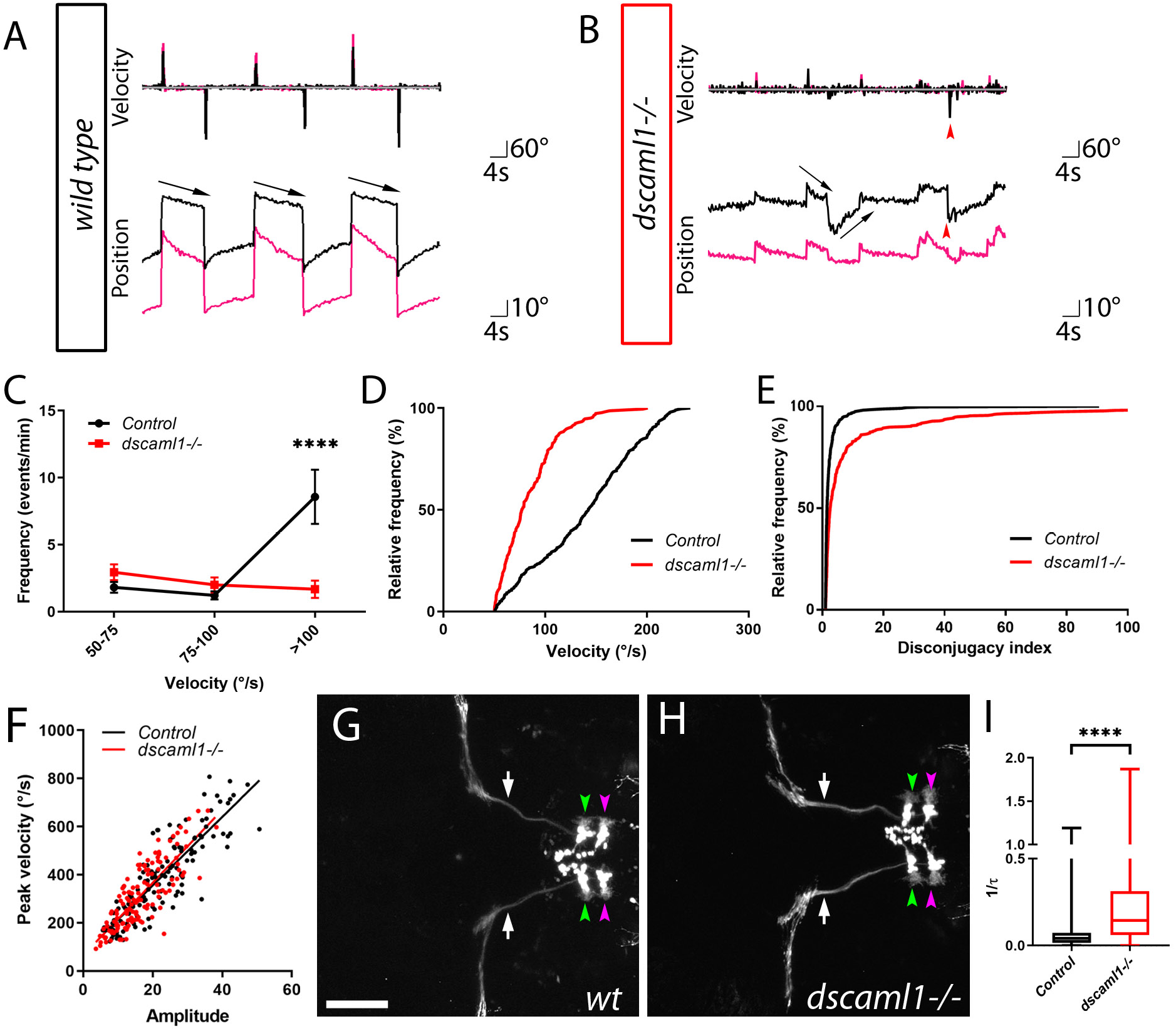
Spontaneous saccade and fixation are abnormal in *dscaml1* mutants. **A-B**, spontaneous saccades in control (A) and *dscaml1−/−* (B) were recorded during uniform illumination (no gratings). Right and left eye traces are in black and fuchsia, respectively. Red arrowheads indicate disconjugated saccades. **C**, the frequency of eye movements divided into angular velocity bins. Controls had significantly more eye movements greater than 100°/s compared to mutants, whereas mutants had more eye movements in the slowest bin compared to control (*p<0.05, ****p<0.0001, two-way ANOVA). D, the relative frequency distribution of eye movements based on angular velocity. Control animals perform significantly more fast eye movements than mutants (p<0.0001, two-sample K-S test). **E**, cumulative distribution of saccade disconjugacy index for controls and mutants (higher equals more disconjugated, see methods). Control index has a median value of 1.56 across the population while the mutant index has a higher median value of 2.33. Distribution is significantly different between control and mutant (p<0.001, two-sample K-S test). **F**, peak velocity and amplitude for each saccade event was plotted, along with linear regression (solid line). The slope is not significantly different between control and mutant (p=0.46). **G-H, an** example of TagRFP-T expressing abducens motor neurons [from *Tg(mnxl:TagRFP-T)]* in wild-type (G) and mutant (H) animal. Dorsal view, rostral to the left. Labeled structures: motor axons (VIth nerve, white arrows), rostral abducens complex (green arrowheads), and caudal abducens complex (magenta arrowheads). Scale bar is 100 μm. **I**, eye position decay (arrows in A, B) calculated as 1/τ. ****: p<0.0001, Mann-Whitney *U* test.

The reduced saccade velocity in mutants may reflect slower eye movements caused by dysfunction in the oculomotor periphery. Alternatively, reduced saccade velocity may be coincidental to smaller saccade amplitudes, which generally have lower velocity. We tested the correlation of peak velocity and saccade amplitude, a linear relationship known as main sequence^19, 41^. Main sequence is a clinically relevant diagnostic metric for saccade: patients with COMA have smaller saccade but normal velocity (normal main sequence), whereas patients with saccade deficits associated with neurodegenerative disorders have both smaller saccade and lower velocity (lower main sequence)^17, 42^. We tested the spontaneous saccade main sequence (sampled at 30 Hz) and found that there was no significant difference between mutants and controls (p=0.468, ANCOVA) (Fig. 6F). In other words, saccades in *dscaml1* had smaller amplitude, but velocity was normal for the given amplitude. The normal main sequence in mutants also suggests that the ocular periphery is not significantly affected by the loss of *dscaml1*. Consistent with this finding, we did not see any gross abnormality in the abducens motor neuron projection to the lateral rectus muscle in the mutants (Fig. 6G, H).

In addition to saccade deficits, post-saccade fixation was significantly perturbed in *dscaml1* mutants, indicating a deficit in the neural integrator pathway. The integrator pathway provides tonic activation of motor neurons to maintain fixation, counteracting spring forces in the eye plant that would cause a drift back to the null position (arrows in Fig. 6A, B); during sustained OKR in a given direction, this pathway is also necessary for converting tonic eye velocity commands to ramplike eye position commands. We modeled this drift rate using an exponential decay function and found that eye position in mutant animals drifted towards baseline more rapidly than in controls (median τ = 24 and 7 seconds for control and mutant animals, respectively, Fig. 6I). Together, these results suggest that *dscaml1* is involved in the function of the saccadic and integrator pathways, but does not affect the function of the oculomotor periphery^43^.

### Loss of *dscaml1* leads to neural activity deficits during OKR fast phase

To understand the neurophysiological basis of the oculomotor phenotypes, we focused on the activity of the abducens motor complex (ABD), which controls the extraocular muscles for horizontal eye movements and serves as the convergence point for different premotor inputs (Fig. 1A)^44,45^. We performed two-photon calcium imaging in control *(dscaml1+/−)* and mutant *(dscaml1−/−)* fish in *elavl3:H2B-GCaMP6f* transgenic background while they performed OKR^46^. Eye positions were recorded simultaneously with an infrared video camera and oculomotor behavior-encoding ABD neurons (motor or internuclear) were then identified based on anatomical location and calcium activity (Fig. 7A-B)^13^. To induce a mixture of optokinetic and spontaneous responses we projected a grating stimulus in front of the animal and then moved that stimulus in a repeating temporal pattern (Fig. 7C-D). We measured mean fluorescence following the onset of stimulus movement in either the ipsilateral (e.g., clockwise for right ABD) or contralateral (e.g., counterclockwise for right ABD) direction. Slow phase ABD activity (a proxy for vestibular pathway activity) was calculated with deconvolved calcium signal for the first three seconds of the ipsilateral or contralateral phases. Fast-phase ABD activity (a proxy for saccadic pathway activity) was calculated with deconvolved average fluorescence following saccadic eye movements in both directions. Since ABD neuron responses are direction selective, we could excite neurons in both hemispheres by alternating the direction of stimulus movement.

**Figure 7.**
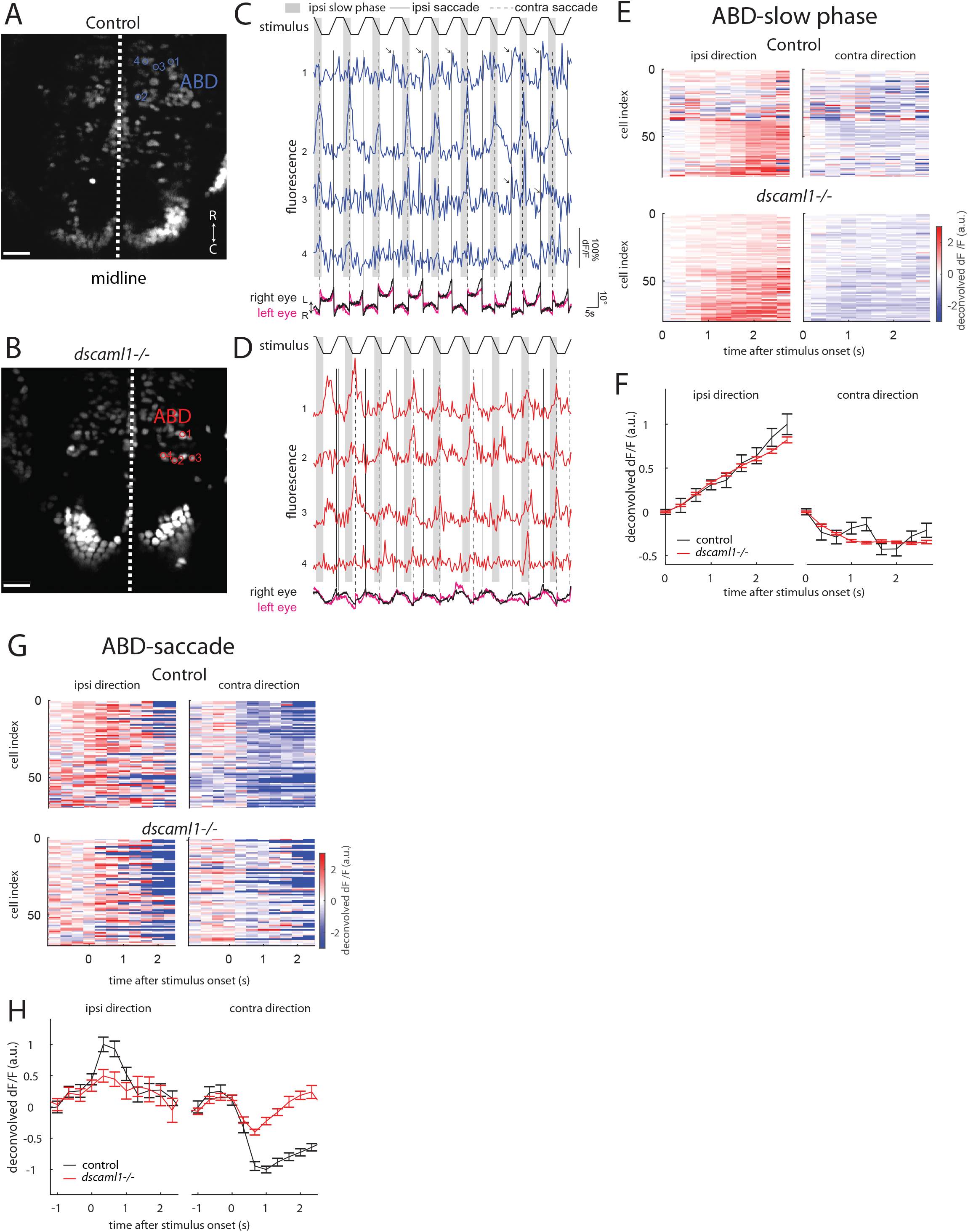
Two-photon calcium imaging in Abducens and Inferior Olive neural populations. **A-D**, two-photon calcium imaging and simultaneous eye position recording during OKR. Circles in time-averaged images show the locations of cells (A, B) with corresponding indices whose fluorescence activity is shown (C, D). Circles in the abducens motor complex (ABD) are marked. E, average fluorescence traces of individual cells (Y-axis) aligned to stimulus onset during slow-phase activity (X-axis) in the ABD. **F**, Population average of fluorescence traces in E. **G**, average fluorescence traces aligned to fast-phase eye movements for cells in the ABD. **H**, activity in G averaged across the population. Error bars in F and H show the median absolute deviation of singlecell activities from the population average divided by the square root of the number of cells, normalized so that population responses are between −1 and 1.

Loss of *dscaml1* did not affect ABD activity during slow phase (n=86 cells from 3 control animals; n=424 cells from 6 mutant animals). Population-average activity in the ABD was comparable in mutant and control animals following the onset of stimulus movement in the ipsilateral direction (Fig. 7E, F). In the contralateral direction, ABD population-average activity reached similar steady-state levels in mutant and controls. This result is consistent with *dscaml1’s* relatively mild effect on slow phase at short time scale and suggests that the vestibular pathway is unaffected.

In contrast, ABD activity was substantially reduced in mutants during fast phase, relative to controls. The saccade-triggered average activity (STA) of ABD cells in the control animals were characterized by a rapid increase or decrease in deconvolved fluorescence following an ipsilateral or contralateral saccade, respectively (Fig. 7G, H). The STA of mutant ABD cells following fast-phase events in either direction was reduced in both directions, compared to controls. Comparing the distribution of trial-averaged STA amplitudes, *dscaml1* mutants had significantly lower amplitudes than controls (p<0.05, two-sample K-S test, n=110 cells from 3 control animals, n=253 cells from 7 mutant animals).

Together, our results show that oculomotor deficits originate centrally and that *dscaml1* affects ABD response selectively during the fast phase. The population-level calcium response in ABD suggests that mutant ABD neurons have normal ramping during the slow phase but lack the saccade-associate burst activity seen in controls. These findings support the idea that *dscaml1* plays a specific role in the function of the saccadic pathway.

## Discussion

We investigated the cellular and behavioral roles of *dscaml1* in zebrafish and took advantage of the oculomotor system to deduce *dscaml1’s* function within a defined neural circuit. Our results underscore the importance of DSCAM proteins in retinal patterning and visuomotor function. Our neurophysiological findings further showed that loss of *dscaml1* leads to impairment in the saccadic but not the pretectum-vestibular premotor pathway, indicating a subcircuit requirement for *dscaml1*. The collection of oculomotor deficits in *dscaml1* mutants bears a striking resemblance to human COMA, for which no animal models exist.

### dscaml1 *and retinal development*

Our results show that zebrafish *dscaml1*, like its mammalian ortholog, is required for maintaining cellular spacing and refining neurites into discrete synaptic laminae in the retina. In the retina, distinct cell types are organized in a mosaic pattern horizontally, and neurites stratify vertically in precise layers in the inner and outer plexiform layers^47^. Both aspects of spatial patterning require the function of DSCAMs. We show that in zebrafish, loss of *dscaml1* causes serotonergic amacrine cells to aggregate. In the IPL, axon terminals of PKCα-positive ON-bipolar cells are more diffuse in *dscaml1* mutants, which likely affects laminar specific synaptogenesis. As ON-bipolar cell axon maturation is activity independent, this diffuse terminal morphology reflects a delay or failure in axon terminal development^26,27^. Interestingly, Dscam rather than Dscaml1 is involved in ON-bipolar cell axon stratification in mice. Nevertheless, Dscaml1 is involved in the refinement of neurite stratification in both mice (VGLUT3+ amacrine cells) and chicken (retinal ganglion cells)^6, 23^. These results demonstrate the remarkable functional conservation of Dscaml1 across vertebrate species.

How the retinal patterning deficits observed in *dscaml1* mutants affect visual function in *dscaml1* still remains to be tested, but the collection of behavioral phenotypes provides some clues. The reduced light-on locomotor response and darker pigmentation in *dscaml1* mutants are suggestive of a reduction in ON pathway function. Consistent with this, the sluggish locomotor response to light-on observed in the *dscaml1* mutant animals resembles the *no optokinetic response c (nrc)* mutant, which completely lacks retinal ON-responses^48^. The more diffuse targeting of ON-bipolar cell axons in *dscaml1* mutants may lead to reduced strength or specificity of light-on responses. However, in contrast to *nrc* animals, which cannot perform OKR, *dscaml1* animals can still perform OKR. Therefore, it is most likely that ON-response is reduced but not completely abolished in *dscaml1* mutants.

### Oculomotor behavior and neural circuit function

The combination of behavioral assays and functional imaging showed that *dscaml1* was necessary for the function of the oculomotor circuit. We focused our analysis to three subcircuits for horizontal eye movements: the saccadic, vestibular, and integrator premotor pathways.

Loss of *dscaml1* strongly affected the saccadic pathway, as deficits were observed in mutants during both reflexive saccades (OKR) and spontaneous scanning saccades. This is consistent with physiological observations showing that saccade-related activity in the abducens was significantly decreased in mutants compared to controls. Retinal defects may contribute to the saccade deficits observed in mutants, at least during reflexive saccades, but would not account for the full extent of the phenotype. Instead, our results favor the hypothesis that abnormal function of the saccadic premotor pathway is the primary cause of the saccade phenotype in *dscaml1* mutants. It is hypothesized that the saccade generator circuit relies on strong local recurrent feedback to generate large and coordinated pulses^49^. This feedback may be easily disrupted through the expected connectivity deficits induced by *dscaml1* mutation. Another factor contributing to the saccadic deficit could be attenuated connectivity at the EBN-ABD synapse. However, the disconjugation of saccades observed in *dscaml1* mutants suggests that the pulse signal from the saccade generators is already fragmented or weakened before being projected to the ABD.

We saw relatively mild deficits in the vestibular pathway. OKR slow phase performance at short time scale was mostly normal in mutants, consistent with the normal ABD calcium dynamics during slow phase. *dcamll* mutant animal’s performance dropped off at higher speeds, but the effects were relatively mild. Given the essential role of the retinal ON pathway in OKR^48^ and the diminished behavioral responses to light onset (locomotor and background adaptation), the normal slow phase performance may be due to compensatory mechanisms that overcome reduced retinal sensory input. It is also worth noting that torsional VOR was affected more strongly than horizontal OKR, indicating that *dscaml1’s* effect on different subcircuits of the oculomotor system is not uniform.

Loss of *dscaml1* resulted in a notable deficit in the function of the integrator pathway. This is most clearly seen in the fast decay of eye fixation after spontaneous saccade in *dscaml1* mutants, relative to controls. Additionally, slow phase deficits in mutants may partly arise from the integrator pathway, which is necessary to integrate the velocity signal from the vestibular nuclei to encode a smooth ramp of eye position^50^. The behavioral dysfunction observed in the mutant could arise from a deficit in the circuit connectivity within the integrator for supporting and coordinating persistent firing^45,51^.

Lastly, the pronounced time-dependent fatigue and lock up phenotypes in *dscaml1* mutants are likely contributed by a combination of different pathways and broader effects beyond the oculomotor circuit. Decreased neurotransmitter release from optic nerve terminals and habituation of retinorecipient neurons have previously been shown to degrade visual response under prolonged stimulation; these mechanisms may underlie the reduced robustness in *dscaml1* mutants^35, 36^. Similarly, lock up may result from changes in neuronal excitability and seizure-like episodes, analogous to what was seen in the zebrafish *didy* mutants (see next section)^34^.

### *Comparison with* didy (*Nav1.1b*) *mutants*

Some aspects of the *dscaml1* phenotype are similar to a previously described zebrafish mutant, *didy*, which encodes the voltage-gated sodium channel Scn1lab (Nav1.1b)^34^. *didy* mutants have spontaneous seizure-like brain activity and very infrequent spontaneous saccades. After 15 seconds of continuous OKR stimulation, *didy* mutants cease to initiate resetting saccades, resulting in lock up of the eyes. Another similarity between *didy* and *dscaml1* is defective light adaptation (darker pigmentation, slow response to light stimulus). *didy* and *dscaml1* may both affect the saccadic pathway, which progressively loses excitability in *didy* mutants. It is important to note, however, that the saccade phenotypes are distinct between *didy* and *dscaml1*. In *didy* animals, saccades have normal speed and amplitude^34^, whereas saccades in *dscaml1* animals are slow and small. Furthermore, lock-up events in *dscaml1* mutants occur more sporadically and across the position range, whereas lock up only occurs at the most eccentric position at the end of a slow phase in *didy* mutants. These distinctions suggest that *didy* and *dscaml1* likely affect saccade generation through independent mechanisms

### Relevance to human oculomotor disorder

*dscaml1* mutant fish share several features of COMA: failure to initiate saccadic eye movements, hypometric saccades, normal main sequence, and intermittent lock up during horizontal OKR. COMA is an infantile-onset condition involving failure of both voluntary and reflexive saccadic eye movements^16, 17,52^. As it is not a true apraxia (where only voluntary movements are affected), the condition is also known as intermittent saccade initiation failure or infantile-onset saccade initiation delay. The etiology of COMA is still poorly understood. Genomic regions surrounding the causative gene for juvenile nephronophthisis *(NPHP1)* have been suggested to contribute to COMA, but mutations in *NPHP1* itself do not consistently cause COMA^53^. *NPHP1* has been linked to primary cilia function, which is crucial for the development of the cerebellum, a key region involved in saccade control^54, 55,56^. Although there is currently no genetic association between human *DSCAML1* and COMA *(DSCAML1* resides on a separate chromosome from *NPHP1)*, further analysis of the *dscaml1* mutants (e.g., cerebellum and extraocular motor neurons) may provide insights to the pathogenesis of COMA and the development of neural circuitry for saccades in general.

### Conclusions

Our investigations on zebrafish *dscaml1* revealed essential roles for a *DSCAM* family gene in visuomotor behavior and subcircuit activity. Given the structural conservation of subcortical circuits and the functional conservation of *dscaml1’s* roles in retinal patterning, it is plausible that the mammalian *DSCAML1* will also contribute to visuomotor processing. By taking advantage of the translucent larval zebrafish system, we recorded oculomotor circuit output dynamics in behaving animals and uncovered a specific dependence of the saccade pathway on *dscaml1*. Our physiological findings in both control and mutant contexts provide a neural basis for the saccade deficits seen in *dscaml1* mutants and potentially for human saccade palsy (COMA). In addition to saccade deficits, our broad examination of oculomotor behaviors also revealed *dscaml1’s* function in VOR, neural integration, and behavioral robustness. These behavioral characterizations provide links between *dscaml1* and diverse aspects of sensorimotor function and will facilitate future studies on the development and disorders of sensorimotor circuits.

## Materials and Methods

### Zebrafish husbandry

Zebrafish (all ages) were raised under 14h light/10h dark cycle at 28.5°C. Embryos and larvae were raised in water containing 0.1% Methylene Blue hydrate (Sigma-Aldrich). At 24 hours postfertilization, embryos used for histological analyses were transferred to E3 buffer containing 0. 003% 1-phenyl-2-thiourea (PTU; Sigma-Aldrich) to prevent pigment formation. Developmental stages are as described by Kimmel et al.^57^. All experimental procedures are performed in accordance with Institutional Animal Care and Use Committee guidelines at Augusta University, Virginia Tech, and Weill Cornell Medical College.

### Mutant and Transgenic Zebrafish lines

*dscaml1* (ZFIN gene name: *Down syndrome cell adhesion molecule like 1)* mutant was generated in TL/AB mixed background using TAL effector nucleases (TALENs) as previously described^21,58^. Two alleles were identified, one harbored a 6 base pair insertion (in frame) and the other harbored a 7 base pair deletion (frame shift). The 7 base deletion mutant (*dscaml1^vt1^*) was used for further analysis. The *dscaml1^vt1^* allele generates a HaeIII (New England Biolabs) restriction site, which was used to distinguish between wild-type and *dscaml1^vt1^* alleles. DNA prep and PCR were performed as described previously^59^, followed by HaeIII digestion for 2 hours at 37°C (primer sequences: aaatactgcacggtgcacacgtc and atgcagatcctacagcctcataatc). After HaeIII digestion, wild-type band was 395 base pairs (uncut) whereas mutant bands were 315 and 75 base pairs. Sequencing of *dscaml1* transcript confirmed incorporation of the 7-base deletion into the open reading frame. *Tg(atoh7:GAP-RFP)* animals were provided by Owen Randlett (Harvard University, Cambridge, MA, USA)^28^. *Tg(elavl3:H2B-GCaMP6f)* and *casper (nac−/−;roy−/−)* animals were provided by Misha Ahrens (HHMI Janelia Farm Research Campus, Ashburn, VA, USA)^46^.

### Image acquisition and processing

Imaging procedures were as previously described^60^. Fluorescent images were acquired using an Olympus FV1000 laser-scanning confocal system with a 20x XLUMPlanFl water-immersion objective or a Nikon A1R MP+ laser scanning confocal system with a CFI75 Apochromat LWD 25x water-immersion objective. Larvae were immobilized with 0.01% tricaine methanesulfonate (MS-222, Sigma-Aldrich), embedded in molten 1.5% low-melt agarose (Fisher Scientific) in a glass-bottomed Petri dish (P50G-1.5-14-F, MatTek). Fish were mounted so that the surface to be imaged was facing the glass bottom.

Images were processed with Fiji^61^ and Photoshop (Adobe Systems) software. To measure the extent of serotonergic amacrine cell aggregation, we acquired confocal stacks and determined the center of mass for each 5-HT positive cell in (x, y, z) coordinates. Intercellular distance between two cells was calculated by the distance formula: 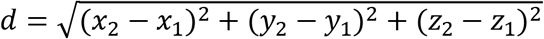. For each cell, intercellular distance to all other cells in the same retina were calculated. Cells with neighboring cells within 3 cell diameter (≤10 microns) were categorized as aggregated. For each eye, the number of aggregating cells was divided by the total number of cells to calculate aggregation ratio.

Quantification of the distribution of PKCα immunolabeling was performed as described by Nevin *et al*.^27^ Using Fiji, a rectangle was drawn across a relatively flat section of the IPL so that the top and bottom of the rectangle abut the boundaries of the IPL, as defined by SV2 immunolabeling. Next, the Plot Profile function was used to measure the average fluorescence intensity across the thickness of the IPL. After exporting numerical data to Excel (Microsoft Inc.), data were normalized to maximal fluorescence intensity and relative position within the IPL. The number of serotonergic neurons were estimated by manually counting 5HT+ cells from 7 optical sections (10μm apart, at horizontal levels adjacent to the lens) in each fish.

For 3D rendering of retinal afferents, *Tg(atoh7:GAP-RFP)* transgenic fish were fixed with 4% PFA and mounted laterally. The eye on the imaged side was removed to allow visualization of the optic tract. 3D rendering was created using Nikon NIS-Elements software. Measurements of images were analyzed using the Prism 6 statistic software (GraphPad).

### *Fluorescent* in situ *hybridization and immunohistochemistry*

*dscaml1* probe was generated by 5’RACE (Smart RACE cDNA Amplification Kit, Clonetech) using 3’ primers designed from Ensembl exon predictions. Amplified DNA was cloned into the pCRII-TOPO vector by TA cloning (Invitrogen). Probe sequence includes 127 base pairs of 5’ untranslated region and 545 base pairs of coding sequence. DIG-labeled *dscaml1* probe synthesis and whole mount *in situ* hybridization were performed as previously described^60^. For 3 and 5 days post-fertilization (dpf) samples, Dextran sulfate (Sigma-Aldrich) and 4-iodophenol (Fluka) were added to the hybridization and tyramide solution to increase signal intensity^62^. Whole-mount immunohistochemistry was performed as described by Randlett et al.^63^. Primary antibodies used were: Znp-1 (anti-synaptotagmin2, Developmental Studies Hybridoma Bank), anti-acetylated-tubulin (Sigma), anti-SV2 (Developmental Studies Hybridoma Bank), anti-HuC/D (Invitrogen), anti-HNK-1 (zn-12, Developmental Studies Hybridoma Bank), anti-5-HT (Sigma), anti-PKCα (Santa Cruz Biotechnologies), zpr-1 (Zebrafish International Resource Center), and anti-Blbp (Abcam). Alexa fluor-conjugated secondary antibodies were used after primary antibody incubation.

### Locomotor assay

Individual 5 dpf larvae were placed into each well of a 24-well tissue culture plate (Fisher Scientific) and transferred into a Zebrabox imaging chamber (Viewpoint). Locomotor activity of each larva was tracked over 24 hours, with white LED illumination turned off at 10 pm and on at 8 am. Total displacement over time was integrated every 10 minutes, measured as previously described^31^.

### OKR and saccade assays

VisioTracker 302060 (New Behavior TSE) was used for OKR and saccade assays. Eye movements of individual fish were recorded by an overhead CCD camera. Zebrafish larvae were placed in the center of a 50mm glass bottom petri dish (MatTek) and immobilized in 1.5-2% low melting agarose (Fisher Scientific) in E3 buffer. Agarose around the eye was removed to allow free eye movement. The dish was then filled with E3. To test slow phase performance under short periodicity, the direction of black and white grating switched every 3 seconds with grating velocity at 7.5°/s. Each experimental run (trial) was 108 seconds long and included twelve 9-second phases at varying contrast levels (0.99, 1.0, 0.5, 0.2, 0.1, 0.05, 0.02, 0.05, 0.1, 0.2, 0.5, 1.0). For each animal, 5-6 trials were tested and OKR response typically initiated during the first three trials (initiation trials). Contrast sensitivity was calculated using trials recorded after the initiation trials. To exclude the effect of response latency and the initial ramping up of eye velocity at stimulus onset, the first 1 second of each 3-second half period was excluded from analysis^33^. Spontaneous saccades and OKR performance under long periodicity were tested using a trial that contains four phases: (1) uniform illumination (1-160s), (2) square wave grating with direction switching every 40s, contrast=1, spatial frequency=0.05 cycles/°, and velocity=10°/s (161-400s), (3) uniform illumination (401480s), (4) square wave grating with direction switching every 8s, contrast=1, spatial frequency=0.05cycles/°, and velocity=10°/s (481-560s).

A saccadic event was defined as sample periods where the instantaneous absolute eye velocity is greater than 50°/s. Slow-phase eye velocity was measured as the mean, saccade-removed, instantaneous velocity averaged across time within 1-second bins. Instantaneous velocity was measured as the difference in smoothed eye position divided by the sample period (200ms, 5 Hz). Eye position was smoothed using a median filter (Matlab medfilt1) of 5 sample periods (1s). We excluded sample periods that fell between 0.5 seconds before and 2 seconds after each saccade to remove the effects of saccade filtering and post-saccade plant relaxation^64^. To analyze lock-up probability and saccade conjugacy we measured eye velocity independently for each eye since lockup could occur independently on a given eye. Lock up periods are defined as 1-second bins where average velocity is <1°/s. Lock up probability was defined as the average number of lock up periods across trials and both eyes. We defined a disconjugacy index as the ratio of left and right instantaneous eye velocity magnitude at each time bin where a saccadic event occurs. The index is defined so that the larger saccade velocity is in the numerator which means the index is always greater than or equal to 1. For all other plots, instantaneous velocity was averaged across both eyes before binning.

To calculate the main sequence of saccade, eye positions were recorded at 30 Hz (33 ms sample period). Saccade peak velocity and amplitude were measured as described by Chen et al.^41^ The main sequence was calculated as the slope of the linear regression for peak velocity and amplitude, using the Prism6 software (Graphpad)^19^.

To measure the drift rate in eye position following spontaneous saccades during the uniform illumination period of the stimulus, we used a quasi-Newton unconstrained optimization (Matlab fminunc with Algorithm set to quasi-Newton) to minimize the squared error between eye position and the function

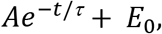

with variable parameters A and *τ*. *E*_0_ was fixed to the mean position for the eye being fit. The algorithm was initialized with tau set to 10 seconds and A set to the first eye position value in the sample. We excluded positions that occurred between 0-1 seconds after each saccade, to avoid fitting post-saccadic relaxation related to plant mechanics. Since the duration of spontaneous fixations was variable, the fixation window analyzed was variable with typical values between 5-20 seconds. We only analyzed exponential fits that passed the following goodness-of-fit criteria: sum-of-squared errors was less than (30°)^2^ or one minus the ratio of mean squared-error to the sample variance was greater than 0.4 and the sum-of-squared errors was less than (200°)^2^. These criteria were chosen based on visual inspection of fit qualities.

### VOR assay

Torsional eye movements were measured in 5 days post-fertilization fish in response to step tilts delivered using an apparatus similar in design to Schoppik *et al*. 2017^38^. All experiments took place in the dark. Larval fish were immobilized completely in 2% low-melting-temperature agar (Thermo Fisher), and the left eye freed. The agar was then pinned (0.1mm stainless minutien pins, FST) to a ~5mm^2^ piece of Sylgard 184 (Dow Corning) which was itself pinned to Sylgard 184 at the bottom of a 10 mm^2^ optical glass cuvette (Azzota). The cuvette was filled with 1ml of E3 media and placed in a custom holder on a 5-axis (X, Y, Z, pitch, roll) manipulator (ThorLabs MT3 and GN2). The fish was aligned with the optical axes of two orthogonally placed cameras such that both the left utricle and two eyes were level with the horizon (front camera). The experimenter running behavior was blind as to the genotype of the fish.

The eye-monitoring camera (Guppy Pro 2 F-031, Allied Vision Technologies) used a 5x objective (Olympus MPLN, 0.1 NA) and custom image-forming optics to create a 100×100 pixel image of the left eye of the fish (6μm/pixel), acquired at 200Hz. The image was processed on-line by custom pattern matching software to derive an estimate of torsional angle (LabView, National Instruments), and data were analyzed using custom MATLAB scripts. A stepper motor (Oriental Motors AR98MA-N5-3) was used to rotate the platform holding the cameras and fish. The platform velocity and acceleration were measured using integrated circuits (IDG500, Invensense and ADXL335, Analog Devices) mounted together on a breakout board (Sparkfun SEN-09268). Fish were rotated stepwise for 4 cycles: from 0° to −15°, where positive values are nose-down, then from −15° to 0°, from 0° to 15°, then back to 0°. Steps had a peak velocity at 35°/sec. The inter-step interval was 7.5 seconds.

The eye’s response across the experiment was first centered to remove any offset introduced by the pattern-matching algorithm. Data were then interpolated with a cubic spline interpolation to correct for occasional transient slowdowns (i.e., missed frames) introduced by the pattern-matching algorithm. The eye’s velocity was estimated by differentiating the position trace; high-frequency noise was minimized using a 4-pole low-pass Butterworth filter (cutoff = 3Hz). Each step response was evaluated manually; trials with rapid deviations in eye position indicative of horizontal saccades or gross failure of the pattern-matching algorithm were excluded from analysis. Across all fish and all steps used to measure the behavior, the median number of usable responses was 7/10. The response to each step for a given fish was defined as the mean across all responses to that step across cycles. The gain was estimated by measuring the peak eye velocity occurring over the period 375-1000ms after the start of the step. Only steps away from the horizon were analyzed.

Of 9 fish, one was excluded because it had fewer than ten steps for analysis, all others had at least ten. The median number of steps ± interquartile range was 18/15±13.75/10.5 for nose-down/nose-up steps respectively.

### Two-Photon Calcium Imaging during Behavior

Embryos from crosses of *dscaml1* heterozygous mutants, *(dscaml1+/−;casper+/−* X *dscaml1+/−; nac+/−; elavl3:H2B-GCaMP6f)* were used. At 5-7 dpf, pigmentless *(nac−/−) Dscaml1* heterozygous and homozygous mutant siblings were immobilized in a gel of 1.8% low-melting temperature agarose (Sigma-Aldrich) in preparation for imaging. Agarose was removed from the eyes to allow them to move freely during imaging. Each fish was genotyped following imaging. Simultaneous eye tracking and two-photon calcium imaging were performed using a custom-built system as previously described^13^. Each image was acquired by raster scanning a mode-locked excitation laser (wavelength set to 930 nanometers) through a 40x water immersion lens to a horizontal plane at the abducens motor neurons. The laser power at the sample varied between 15-25 mW. For each animal, we recorded 3-10 planes at a rate of 1.95Hz and duration of 5 minutes per plane. Each plane was recorded while vertical stripes were projected onto a screen of diffusion film placed 1-3 cm in front of the animal providing an optokinetic stimulus^13^. The stripes moved at a constant velocity whose magnitude and direction changed in a repeating pattern that consisted of equal durations of positive, negative, and zero velocity, with each phase lasting 3.14 (2 heterozygotes and 6 mutants) or 31.26 seconds (1 heterozygotes and 1 mutant). Eye position was measured using a sub-stage, infrared camera (Allied Vision Technologies, Guppy FireWire camera) that acquired frames at 13 Hz^13^.

### Identification of putative abducens neurons

Putative abducens neurons were identified based on cell location and fluorescence activity. Abducens motor neurons are clustered in rhombomeres (rh) 5-6 and are arranged in dorsal-ventral columns. The imaging window (185 μm^2^) was positioned over rh 5-6 using the posterior otolith as guides to image the abducens population^65^. We only examined cells whose fluorescence responses were correlated with direction-rectified eye-position and/or eye-velocity traces with an absolute Pearson correlation coefficient greater than or equal to 0.3^13, 44,51^. Before correlation calculation, we convolved eye-position and eye-velocity variables with a 2 second exponentially decaying calcium impulse response function to account for the calcium buffering associated with a cell’s action potential^44,66^. Single neurons that contained at least one pixel with an absolute correlation value above 0.3 were manually selected for further analysis. To correct for animal motion artifacts, fluorescence movies were first pre-processed using a procedure that relies on cross-correlation of individual frames with a time-averaged reference frame^51^. Frames that undergo large shifts from the reference frame (greater than the median plus 5 times the median absolute deviation) were excluded from analysis. To compute delta F over F time series (dF/F), we subtracted and then divided the time-averaged fluorescence within each ROI.

### Saccade and Stimulus-Triggered Average Calculation

For each cell, we computed the average dF/F response relative to saccadic (fast-phase) eye movements and relative to the start of optokinetic stimulus movements. A fast-phase event is defined as described above (50 °/s). Eye velocity is computed as the instantaneous difference in smoothed eye position divided by instantaneous eye sampling time (77 ms). Eye position was smoothed using a median filter (Matlab medfilt1) of order equivalent to 500 ms. In order to combine dF/F traces across saccadic events and the start of stimuli, we linearly interpolated saccade or stimulus-aligned responses to a grid of evenly spaced time bins 333 ms in width before averaging. We excluded planes from analysis if there are less than 5 saccades events available for computing the STA. When averaging stimulus-aligned responses we only used traces where a fast-phase event did not occur within the first 3 seconds following stimulus onset (n=3 hets, n=6 mutants).

## Acknowledgments

This work was supported by funding from the National Institutes of Health (R01 EY024844 to Y.A.P., K99 EY027017 to A.D.R., R01 EY027036 to E.R.F.A.), the Medical College of Georgia, and Virginia Tech. We thank the animal care staff at Augusta University and Virginia Tech for animal husbandry, the Augusta University Electron Microscopy and Histology Core for retina histology preparation, A. Pauli for help generating the *dscaml1* TALEN mutants, J. Mathias for assistance with locomotor behavior analysis, F. Ali for technical assistance, and K. Bollinger and J. Sanes for helpful discussions.

## Author Contributions

T.W., M.M., and Y.A.P. conceived the study, with input from A.F.S. Y.A.P., J.A.G. and S.Z. generated the *dscaml1* mutant line, using TALEN targeting constructs designed and made by D.R., S.Q.T., and J.K.J. T.W., M.M., R.R., C.K., and Y.A.P. contributed to the histological analyses. M.M., A.R. A.S., K.E.H., D.S., E. R.F.A. and Y.A.P. contributed to the behavioral analyses. A.R. and E.R.F.A. designed and interpreted the functional imaging experiments. A.R. performed and analyzed the functional imaging experiments with help from M.M. Y.A.P. wrote the manuscript, with contributions from T.W., M.M., A.R., R.R., D.S., S.L.G., and E.R.F.A.

## Figures and Movie

**Movie 1. OKR response**. Control (left) and *dscaml1−/−* (right) animals performing OKR.

**Supplementary Figure S1.**
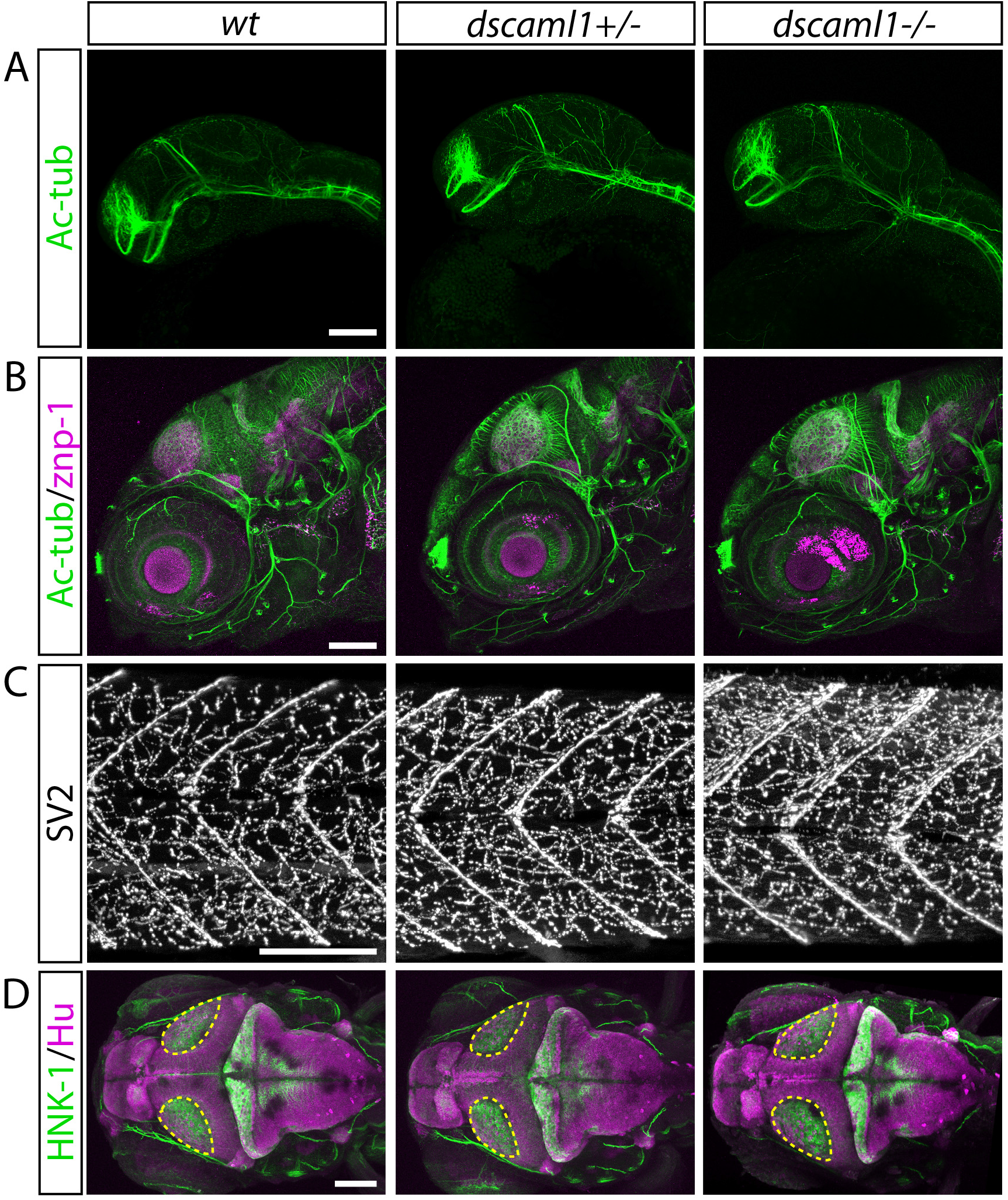
Brain morphogenesis is grossly normal in *dscaml1* mutants. **A-B**, Lateral view of axon tracts (Ac-tub, green) and synapses (znp-1, magenta) at 1 and 5 dpf. In 1 dpf embryos (A), no differences were seen in the formation of the major commissures and longitudinal tracts^67^. In 5 dpf larvae, no apparent abnormalities were seen in the morphology of sensory nerves, motor nerves, or distribution of synapses in the brain and retina^68, 69^. **C**, Lateral view of the neuromuscular junction of the trunk, stained with a marker for presynaptic terminals (SV2). **D**. Dorsal view of 5 dpf larvae stained for mature neurons (Hu, magenta) and axon tracts (HNK-1, green). Areas outlined in yellow are the optic tectum neuropil region. Images show maximal intensity projection of confocal image stacks. Scale bars are 100μm.

**Supplementary Figure S2.**
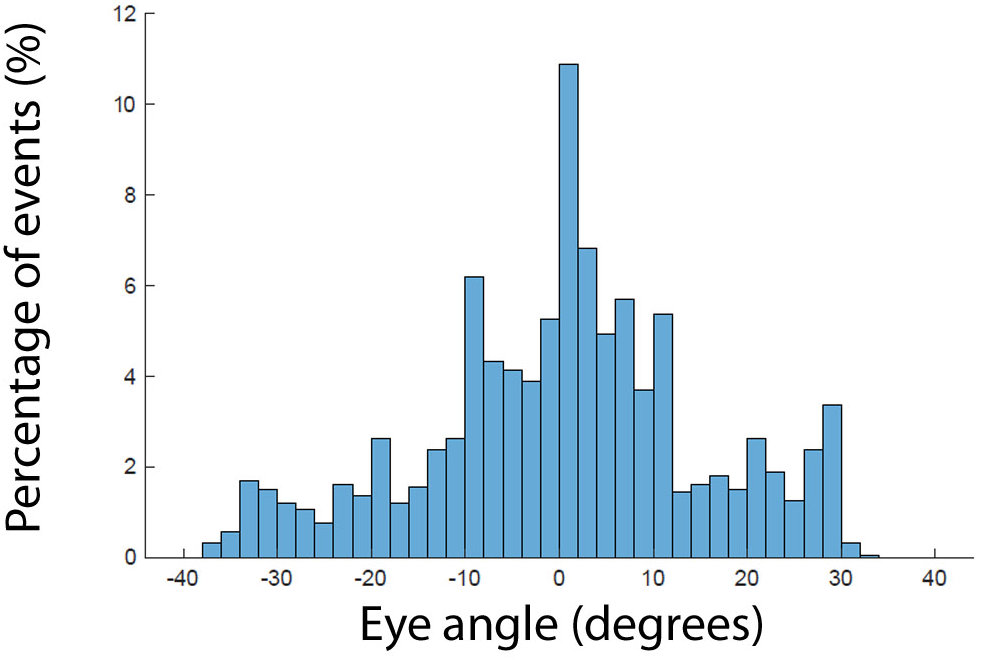
Eye Position during lock-up in *dscaml1* mutants. Histogram of left and right eye positions averaged during 1 second bins where lock-up occurs. Bin size is 2 degrees.

## References

1. Iossifov I, et al. The contribution of de novo coding mutations to autism spectrum disorder. Nature 515, 216–221 (2014).

2. Karaca E, et al. Genes that Affect Brain Structure and Function Identified by Rare Variant Analyses of Mendelian Neurologic Disease. Neuron 88, 499–513 (2015).

3. Blank M, et al. The Down syndrome critical region regulates retinogeniculate refinement. J Neurosci 31, 5764–5776 (2011).

4. Cui S, Lao L, Duan J, Jin G, Hou X. Tyrosine phosphorylation is essential for DSCAML1 to promote dendrite arborization of mouse cortical neurons. Neurosci Lett 555, 193–197 (2013).

5. Maynard KR, Stein E. DSCAM contributes to dendrite arborization and spine formation in the developing cerebral cortex. J Neurosci 32, 16637–16650 (2012).

6. Yamagata M, Sanes JR. Dscam and Sidekick proteins direct lamina-specific synaptic connections in vertebrate retina. Nature 451, 465–469 (2008).

7. Zhang L, Huang Y, Chen JY, Ding YQ, Song NN. DSCAM and DSCAML1 regulate the radial migration and callosal projection in developing cerebral cortex. Brain Res 1594, 61–70 (2015).

8. Bruce FM, Brown S, Smith JN, Fuerst PG, Erskine L. DSCAM promotes axon fasciculation and growth in the developing optic pathway. Proceedings of the National Academy of Sciences of the United States of America 114, 1702–1707 (2017).

9. Leigh RJ, Zee DS. The neurology of eye movements, 5th edition. edn. Oxford University Press (2015).

10. Bittencourt J, et al. Saccadic eye movement applications for psychiatric disorders. Neuropsychiatr Dis Treat 9, 1393–1409 (2013).

11. Anderson TJ, MacAskill MR. Eye movements in patients with neurodegenerative disorders. Nature Reviews Neurology 9, 74 (2013).

12. Galicia CA, Sukeena JM, Stenkamp DL, Fuerst PG. Expression patterns of dscam and sdk gene paralogs in developing zebrafish retina. Molecular vision 24, 443–458 (2018).

13. Daie K, Goldman MS, Aksay ER. Spatial patterns of persistent neural activity vary with the behavioral context of short-term memory. Neuron 85, 847–860 (2015).

14. Straka H, Beck JC, Pastor AM, Baker R. Morphology and physiology of the cerebellar vestibulolateral lobe pathways linked to oculomotor function in the goldfish. J Neurophysiol 96, 1963–1980 (2006).

15. Masseck OA, Hoffmann KP. Comparative neurobiology of the optokinetic reflex. Ann N Y Acad Sci 1164, 430–439 (2009).

16. Harris CM, Shawkat F, Russell-Eggitt I, Wilson J, Taylor D. Intermittent horizontal saccade failure (‘ocular motor apraxia’) in children. Br J Ophthalmol 80, 151–158 (1996).

17. Zee DS, Yee RD, Singer HS. Congenital ocular motor apraxia. Brain 100, 581–599 (1977).

18. Easter SS, Jr., Nicola GN. The development of eye movements in the zebrafish (Danio rerio). Dev Psychobiol 31, 267–276 (1997).

19. Beck JC, Gilland E, Tank DW, Baker R. Quantifying the ontogeny of optokinetic and vestibuloocular behaviors in zebrafish, medaka, and goldfish. J Neurophysiol 92, 3546–3561 (2004).

20. Fuerst PG, et al. DSCAM and DSCAML1 function in self-avoidance in multiple cell types in the developing mouse retina. Neuron 64, 484–497 (2009).

21. Reyon D, Tsai SQ, Khayter C, Foden JA, Sander JD, Joung JK. FLASH assembly of TALENs for high-throughput genome editing. Nat Biotechnol 30, 460–465 (2012).

22. Wagle M, Mathur P, Guo S. Corticotropin-releasing factor critical for zebrafish camouflage behavior is regulated by light and sensitive to ethanol. JNeurosci 31, 214–224 (2011).

23. Garrett AM, Tadenev AL, Hammond YT, Fuerst PG, Burgess RW. Replacing the PDZ-interacting C-termini of DSCAM and DSCAML1 with epitope tags causes different phenotypic severity in different cell populations. eLife 5, (2016).

24. Maurer CM, Schonthaler HB, Mueller KP, Neuhauss SC. Distinct retinal deficits in a zebrafish pyruvate dehydrogenase-deficient mutant. J Neurosci 30, 11962–11972 (2010).

25. Wan L, Almers W, Chen W. Two ribeye genes in teleosts: the role of Ribeye in ribbon formation and bipolar cell development. J Neurosci 25, 941–949 (2005).

26. Schroeter EH, Wong RO, Gregg RG. In vivo development of retinal ON-bipolar cell axonal terminals visualized in nyx::MYFP transgenic zebrafish. Vis Neurosci 23, 833–843 (2006).

27. Nevin LM, Taylor MR, Baier H. Hardwiring of fine synaptic layers in the zebrafish visual pathway. Neural Dev 3, 36 (2008).

28. Zolessi FR, Poggi L, Wilkinson CJ, Chien CB, Harris WA. Polarization and orientation of retinal ganglion cells in vivo. Neural Dev 1, 2 (2006).

29. Robles E, Laurell E, Baier H. The retinal projectome reveals brain-area-specific visual representations generated by ganglion cell diversity. Curr Biol 24, 2085–2096 (2014).

30. Prober DA, Rihel J, Onah AA, Sung RJ, Schier AF. Hypocretin/orexin overexpression induces an insomnia-like phenotype in zebrafish. J Neurosci 26, 13400–13410 (2006).

31. Farrell TC, Cario CL, Milanese C, Vogt A, Jeong JH, Burton EA. Evaluation of spontaneous propulsive movement as a screening tool to detect rescue of Parkinsonism phenotypes in zebrafish models. Neurobiol Dis 44, 9–18 (2011).

32. Kubo F, Hablitzel B, Dal Maschio M, Driever W, Baier H, Arrenberg AB. Functional architecture of an optic flow-responsive area that drives horizontal eye movements in zebrafish. Neuron 81, 1344–1359 (2014).

33. Rinner O, Rick JM, Neuhauss SC. Contrast sensitivity, spatial and temporal tuning of the larval zebrafish optokinetic response. Invest Ophthalmol Vis Sci 46, 137–142 (2005).

34. Schoonheim PJ, Arrenberg AB, Del Bene F, Baier H. Optogenetic localization and genetic perturbation of saccade-generating neurons in zebrafish. J Neurosci 30, 7111–7120 (2010).

35. Smear MC, et al. Vesicular glutamate transport at a central synapse limits the acuity of visual perception in zebrafish. Neuron 53, 65–77 (2007).

36. Perez-Schuster V, et al. Sustained Rhythmic Brain Activity Underlies Visual Motion Perception in Zebrafish. Cell Rep 17, 1098–1112 (2016).

37. Chen CC, et al. Velocity storage mechanism in zebrafish larvae. The Journal of Physiology 592, 203–214 (2014).

38. Schoppik D, et al. Gaze-stabilizing central vestibular neurons project asymmetrically to extraocular motoneuron pools. The Journal of Neuroscience, (2017).

39. Bianco I, Kampff A, Engert F. Prey Capture Behavior Evoked by Simple Visual Stimuli in Larval Zebrafish. Frontiers in Systems Neuroscience 5, (2011).

40. Schmitt EA, Dowling JE. Early retinal development in the zebrafish, Danio rerio: light and electron microscopic analyses. J Comp Neurol 404, 515–536 (1999).

41. Chen CC, Bockisch CJ, Straumann D, Huang MY. Saccadic and Postsaccadic Disconjugacy in Zebrafish Larvae Suggests Independent Eye Movement Control. Front Syst Neurosci 10, 80 (2016).

42. Garbutt S, Harwood MR, Harris CM. Comparison of the main sequence of reflexive saccades and the quick phases of optokinetic nystagmus. Br J Ophthalmol 85, 1477–1483 (2001).

43. Sparks DL. The brainstem control of saccadic eye movements. Nature reviews Neuroscience 3, 952–964 (2002).

44. Miri A, Daie K, Burdine RD, Aksay E, Tank DW. Regression-based identification of behavior-encoding neurons during large-scale optical imaging of neural activity at cellular resolution. J Neurophysiol 105, 964–980 (2011).

45. Vishwanathan A, Daie K, Ramirez AD, Lichtman JW, Aksay ERF, Seung HS. Electron Microscopic Reconstruction of Functionally Identified Cells in a Neural Integrator. Curr Biol 27, 2137–2147 e2133 (2017).

46. Vladimirov N, et al. Light-sheet functional imaging in fictively behaving zebrafish. Nat Methods 11, 883–884 (2014).

47. Hoon M, Okawa H, Della Santina L, Wong RO. Functional architecture of the retina: development and disease. Progress in retinal and eye research 42, 44–84 (2014).

48. Emran F, Rihel J, Adolph AR, Wong KY, Kraves S, Dowling JE. OFF ganglion cells cannot drive the optokinetic reflex in zebrafish. Proceedings of the National Academy of Sciences of the United States of America 104, 19126–19131 (2007).

49. Lo CC, Wang XJ. Cortico-basal ganglia circuit mechanism for a decision threshold in reaction time tasks. Nat Neurosci 9, 956–963 (2006).

50. Cannon SC, Robinson DA. Loss of the neural integrator of the oculomotor system from brain stem lesions in monkey. J Neurophysiol 57, 1383–1409 (1987).

51. Lee MM, Arrenberg AB, Aksay ER. A structural and genotypic scaffold underlying temporal integration. J Neurosci 35, 7903–7920 (2015).

52. Salman MS. Infantile-onset saccade initiation delay (congenital ocular motor apraxia). Curr Neurol Neurosci Rep 15, 24 (2015).

53. Betz R, et al. Children with ocular motor apraxia type Cogan carry deletions in the gene (NPHP1) for juvenile nephronophthisis. J Pediatr 136, 828–831 (2000).

54. Jauregui AR, Nguyen KCQ, Hall DH, Barr MM. The <em>Caenorhabditis elegans</em> nephrocystins act as global modifiers of cilium structure. The Journal of Cell Biology 180, 973–988 (2008).

55. Louie CM, Gleeson JG. Genetic basis of Joubert syndrome and related disorders of cerebellar development. Human molecular genetics 14 Spec No. 2, R235–242 (2005).

56. Matsui H, Namikawa K, Babaryka A, Koster RW. Functional regionalization of the teleost cerebellum analyzed in vivo. Proceedings of the National Academy of Sciences of the United States of America 111, 11846–11851 (2014).

57. Kimmel CB, Ballard WW, Kimmel SR, Ullmann B, Schilling TF. Stages of embryonic development of the zebrafish. DevDyn 203, 253–310 (1995).

58. Sander JD, et al. Targeted gene disruption in somatic zebrafish cells using engineered TALENs. Nat Biotechnol 29, 697–698 (2011).

59. Pan YA, et al. Zebrabow: multispectral cell labeling for cell tracing and lineage analysis in zebrafish. Development 140, 2835–2846 (2013).

60. Pan YA, Choy M, Prober DA, Schier AF. Robo2 determines subtype-specific axonal projections of trigeminal sensory neurons. Development 139, 591–600 (2012).

61. Schindelin J, et al. Fiji: an open-source platform for biological-image analysis. Nat Methods 9, 676–682 (2012).

62. Lauter G, Soll I, Hauptmann G. Multicolor fluorescent in situ hybridization to define abutting and overlapping gene expression in the embryonic zebrafish brain. Neural Dev 6, 10 (2011).

63. Randlett O, et al. Whole-brain activity mapping onto a zebrafish brain atlas. Nature Methods 12, 1039–1046 (2015).

64. Sklavos S, Dimitrova DM, Goldberg SJ, Porrill J, Dean P. Long time-constant behavior of the oculomotor plant in barbiturate-anesthetized primate. J Neurophysiol 95, 774–782 (2006).

65. Miri A, Daie K, Arrenberg AB, Baier H, Aksay E, Tank DW. Spatial gradients and multidimensional dynamics in a neural integrator circuit. Nat Neurosci 14, 1150–1159 (2011).

66. Kawashima T, Zwart MF, Yang CT, Mensh BD, Ahrens MB. The Serotonergic System Tracks the Outcomes of Actions to Mediate Short-Term Motor Learning. Cell 167, 933–946 e920 (2016).

67. Ross LS, Parrett T, Easter SS, Jr. Axonogenesis and morphogenesis in the embryonic zebrafish brain. J Neurosci 12, 467–482 (1992).

68. Fox MA, Sanes JR. Synaptotagmin I and II are present in distinct subsets of central synapses. J Comp Neurol 503, 280–296 (2007).

69. Higashijima S, Hotta Y, Okamoto H. Visualization of cranial motor neurons in live transgenic zebrafish expressing green fluorescent protein under the control of the islet-1 promoter/enhancer. J Neurosci 20, 206–218 (2000).

